# The Mitochondrial Permeability Transition Phenomenon Elucidated by Cryo-EM Reveals the Genuine Impact of Calcium Overload on Mitochondrial Structure and Function

**DOI:** 10.1101/2020.03.13.991349

**Authors:** Jasiel O. Strubbe-Rivera, Jason R. Schrad, Evgeny V. Pavlov, James F. Conway, Kristin N. Parent, Jason N. Bazil

**Affiliations:** Pharmacology and Toxicology, Michigan State University, East Lansing, MI; Biochemistry and Molecular Biology, Michigan State University, East Lansing, MI; Basic Science and Craniofacial Biology, New York University, New York, NY; Structural Biology, University of Pittsburgh School of Medicine, Pittsburgh, PA; Physiology, Michigan State University, East Lansing, MI

## Abstract

Mitochondria have a remarkable ability to uptake and store massive amounts of calcium. However, the consequences of massive calcium accumulation remain enigmatic. In the present study, we analyzed a series of time-course experiments to identify the sequence of events that occur in a population of guinea pig cardiac mitochondria exposed to excessive calcium overload that cause mitochondrial permeability transition (MPT). By analyzing coincident structural and functional data, we determined that excessive calcium overload is associated with large calcium phosphate granules and inner membrane fragmentation, which explains the extent of mitochondrial dysfunction. This data also reveals a novel mechanism for cyclosporin A, an inhibitor of MPT, in which it preserves inner membrane architecture despite the presence of massive calcium phosphate granules in the matrix. Overall, these findings establish a mechanism of calcium-induced mitochondrial dysfunction and the impact of calcium regulation on mitochondrial structure and function.

## Introduction

Mitochondria regulate cell fate through a variety of means (Frank et al., 2001; Halestrap & Pasdois, 2009; Tait & Green, 2013; Youle & Karbowski, 2005; Youle & Narendra, 2011). Their extensive networks and dynamic architecture facilitate metabolic signaling to ensure proper cellular function and survival. Mitochondria achieve this by integrating intracellular cues and physiological stimuli to regulate ATP production, metabolite oxidation, calcium signaling, phospholipid and steroid hormone biosynthesis, and mitochondrial fission and fusion processes (Balaban, 2002; Eisner, Picard, & Hajnoczky, 2018; Friedman & Nunnari, 2014; Glancy, Willis, Chess, & Balaban, 2013; Mannella, Lederer, & Jafri, 2013; Marchi, Patergnani, & Pinton, 2014; Pfanner, Warscheid, & Wiedemann, 2019; Wescott, Kao, Lederer, & Boyman, 2019). As such, mitochondria must operate under a range of physiological conditions including transient changes in energy demand, oxidative stress, and moderate calcium overload. For example, in highly metabolic organs such as heart, brain and kidney, their response to these conditions is crucial for cell survival (Fridolfsson et al., 2012). However, in pathological conditions, such as during an ischemia/reperfusion event, mitochondria undergo a phenomenon known as the mitochondrial permeability transition (MPT). MPT is a gateway mechanism of cell death and involves the opening of a non-selective pore that allows small molecules and metabolites up to 1.5 kDa in size to (Du et al., 2008) freely diffuse across the inner mitochondrial membrane (Bernardi, 1999). When the pore is open, the membrane potential is dissipated, there is a loss of respiratory control, ATP is hydrolyzed, and osmotic swelling occurs (Halestrap & Pasdois, 2009). The swelling causes inner membrane unfolding, outer membrane rupture, and eventually release of apoptogenic molecules, including cytochrome c (cyt. c) that ends in cell death.

While the consequences of MPT are well appreciated, the molecular composition of the pore is currently unknown. The MPT phenomenon was first observed nearly seven decades ago when early studies in the mid 1950’s to early 1960’s demonstrated massive mitochondrial swelling under certain conditions (Chappell & Crofts, 1965; Hackenbrock & Caplan, 1969; F. E. Hunter, Jr. et al., 1964; Lehninger & Remmert, 1959; Raaflaub, 1953). These conditions involved calcium overload, high inorganic phosphate concentrations, fatty acids, oxidative stress, and adenine nucleotide pool depletion. Interestingly, acidosis, adenine nucleotides, divalent cations (e.g., Mg^2+^, Mn^2+^, Ba^2+^ and Zn^2+^), and some metabolic cofactors (Bonsi et al., 2006) prevent pore opening. In the late 1970’s, Haworth and Hunter introduced the term ‘permeability transition’ and highlighted two important points: 1) pore opening is triggered by calcium, and 2) it is closed when calcium is removed from the environment (D. R. Hunter & Haworth, 1979). Their results were later confirmed by Crompton et al. (Crompton, Ellinger, & Costi, 1988) who further proposed that the pore may have a protein identity with some physiological role (Bernardi, 1999; Halestrap & Pasdois, 2009; D. R. Hunter & Haworth, 1979; D. R. Hunter, Haworth, & Southard, 1976). Soon after, other pioneering studies demonstrated that this phenomenon was involved in many pathophysiological diseases and conditions such as neurological disorders, aging, response to toxins, cancer, muscular dystrophy, and ischemia-reperfusion injury (Fridolfsson et al., 2012; Jadiya et al., 2019; Lim et al., 2002; Marchi et al., 2013; Millay et al., 2008). Despite the well-known effects of calcium overload on mitochondrial function, the specific details remain a mystery.

As of now, the current dogma of mitochondrial calcium overload is that mitochondrial dysfunction arises from the opening of a calcium-dependent, free radical sensitized, and proteinaceous molecular pore whose molecular identity thus far remains elusive. Unfortunately, efforts to identify the gene products responsible have been a rollercoaster ride of misleading discoveries and dashed hopes (Baines & Gutierrez-Aguilar, 2018; Carroll, He, Ding, Fearnley, & Walker, 2019; Chinopoulos, 2018). Instead of focusing on the pore, we sought to investigate the consequence of excessive calcium overload on a population of isolated mitochondria by analyzing cryo-electron microscopy (cryo-EM) time-course data. This powerful imaging technique was coupled with high-resolution respirometry and spectrofluorimetry to structurally analyze the effect of calcium overload on mitochondrial function. We identified a novel mechanism that links calcium phosphate granule formation to cristae structural changes, inner membrane fragmentation, and ultimately mitochondrial permeabilization. This mechanism is not mutually exclusive with the current dogma as it integrates many past findings in a concise, overarching theoretical framework. However, our new data add exciting therapeutic targets for mitochondrial-protective therapies.

## Methods

### Ethical Approval

This work conformed to the National Institutes of Health’s Guide for the Care and Use of Laboratory Animals and was approved by Michigan State University’s Institutional Animal Care and Use Committee.

### Mitochondria Isolation and Protein Quantification

Cardiac mitochondria were isolated from guinea pig hearts using differential centrifugation as described in Wollenman et al. (Wollenman, Vander Ploeg, Miller, Zhang, & Bazil, 2017) Briefly, Hartley albino guinea pigs weighing 350 – 450 g (4 – 6 weeks) were injected with heparin (500 units/ml) into the intraperitoneal cavity to prevent blood clotting during the cardiac mitochondrial isolation. Before heart removal, the animals were deeply anesthetized with 4 – 5 % isoflurane. Prior to decapitation by guillotine, a noxious stimulus (paw pinch and eyelid reflex) confirmed the animals were fully sedated. After decapitation, a thoracotomy was performed. The heart was then perfused with cold cardioplegia solution and homogenized as described previously (Wollenman et al., 2017). Mitochondrial protein content was quantified using the BIO-RAD Bovine Serum Albumin (BSA) Standard Set Kit and the BCA assay. The mitochondrial suspension was diluted to a working concentration of 40 mg/ml and kept on ice for the duration of the experiment (4 – 8 hours). Substrate stock solutions were neutralized to pH 7.0.

### Mitochondrial Quality Control

The mitochondrial quality was determined using an Oxygraph 2k (Oroboros Instruments Corp., Innsbruck, Austria) under constant stirring. The O2k chambers were loaded with 2 mL respiratory buffer containing 130 mM KCl, 5 mM K_2_HPO_4_, 20 mM MOPS, and 1 mM MgCl_2_, 1 mM EGTA, 0.1 % (w/v) BSA at a pH of 7.1 and 37 °C. All subsequent experiments were done using this buffer and temperature. At 0 mins, 5 mM sodium pyruvate and 1 mM L-malate were added followed by 0.1 mg/ml mitochondria. Here we defined leak state as the rate of oxygen consumption by mitochondria only in the presence of substrates. At 5 mins a bolus of ADP (500 µM) was added to induce maximal ADP-stimulated respiration. Quality was assessed by computing the respiratory control ratio (maximal ADP-stimulated rate divided by the leak rate). Only mitochondria with an RCR value greater than or equal to 16 were used in the experiments.

### Calcium Contamination and Buffer Calcium Measurements

The amount of contaminating calcium present in the respiratory buffer was 4.0 µM ± 0.43 µM which comes from reagent impurities (Wollenman et al., 2017). This was measured using a perfectION™ calcium selective electrode (Mettler Toledo, Columbus, OH). Results were further confirmed using 1 µM calcium fluorescent indicator calcium green 5N (503 nm excitation and 531 nm emission) using an Olis® DM245 spectrofluorimeter (Olis, Inc., Bogart, GA, USA).

### Calcium Effects on Respiration and Oxidative Phosphorylation

Calcium effects on mitochondrial leak and ADP-stimulated respiration were determined by quantifying changes in leak and ADP-stimulated respiration rates after a calcium challenge in the presence or absence of cyclosporin A (CsA). At 0 mins, 5 mM sodium pyruvate, 1 mM L-malate, ± 1 µM CsA, 0.1 mg/ml mitochondria were injected into each 2 mL chamber containing respiratory buffer. At 5 mins, a calcium bolus of either 75 or 100 µM calcium chloride was injected. At 10 mins, 500 µM ADP was added induce maximal ADP-stimulated respiration.

### Mitochondrial Swelling Assay

Mitochondrial swelling was quantified by measuring absorbance at 540 nm using an Olis® DM245 spectrofluorimeter with a dual-beam absorbance module. At 0 mins, 5 mM pyruvate and 1 mM L-malate was added to a polystyrene cuvette with respiration buffer containing ± 1 µM CsA followed by the addition of 0.1 mg/ml mitochondria. At 5 mins, a 75 or 100 µM calcium chloride bolus was added and the absorbance was recorded for a total of 15 mins. The minimum absorbance signal was determined by adding the uncoupler FCCP (1 µM) and the channel forming peptide Alamethicin (10 µg/mg). To normalize the raw traces, we used the minimum absorbance value followed by the absorbance just before the addition of a calcium bolus.

### Calcium Uptake Dynamics

Calcium uptake dynamics were quantified using the fluorescent dye, calcium green 5N (CaGr5N). Fluorescence was measured using an Olis® DM245 spectrofluorimeter. The dye was excited at 506 nm and the emission recorded at 531 nm. To minimize variability in dye concentration, 1 µM CaGr5N was added to 50 mL stocks of respiration buffer as opposed to adding small volumes to the 2 mL assay volume. At 0 mins, ± 1 µM CsA, 5 mM sodium pyruvate and 1 mM L-malate, and 0.1 mg/ml mitochondria were added to a polystyrene cuvette. At 5 mins, a bolus of either 75 or 100 µM calcium chloride was added and the fluorescence was recorded for 15 mins.

### Cryo-EM Sample Vitrification and Imaging

Isolated mitochondria were suspended at a concentration of 0.1 mg/ml in 2 mL respiration buffer with 5 mM sodium pyruvate and 1 mM L-malate. At the collection times indicated, 5 µL samples were pipetted from the mitochondrial suspension and deposited on Quantifoil R2/2 Holey Carbon grids that had been plasma-cleaned for 20 seconds using a Fischione Instruments model 1020 plasma cleaner. Grids were blotted to thin the water layer, and subsequently plunged into liquid ethane at room temperature using a manual plunge-freezing device (Michigan State University Physics Machine Shop). Grids were then transferred and stored in liquid nitrogen until imaging. Data for the 75 µM calcium chloride experiments were collected in the cryo-EM facility at the University of Pittsburgh School of Medicine using an FEI Polara G2 cryo-electron microscope with a field emission gun operating at 300 kV at nominal magnification of 9,400x with a post-column magnification of 1.4x to obtain a ∼12 – 10 Å/pixel resolution. Images were recorded on an FEI Falcon 3 direct electron-detecting camera. Data for the 100 µM calcium chloride experiments were collected in the cryo-EM facility at the University of Pittsburgh School of Medicine using an FEI TF20 cryo-electron microscope with a field emission gun operating at 200 kV. The images were collected using a nominal magnification in the range of 5,000x on a TVIPS XF416 CMOS camera with a post-column magnification of 1.4x to obtain a 22 Å/pixel resolution. At these magnifications, the electron dose (e^−^ /Å^2^) is low enough to avoid significant sample destruction.

### Calcium Phosphate Granules, Posner’s Clusters, and Mitochondrial Structure Quantification

The program EMAN2 (Tang et al., 2007) was used to quantify the total number of granules for each mitochondrion under each condition from TEM images. A total of 1345 individual mitochondrial images were acquired in the presence and absence of CsA for two calcium treatments. For the 75 µM calcium chloride treatment, there were 235 images of control mitochondria and 645 images of CsA-treated mitochondria. For the 100 µM calcium chloride treatment, there were 231 images of control mitochondria and 234 images for CsA-treated mitochondria. Mitochondrial and phosphate granule diameters were computed from three averages of two diagonal and one horizontal diameter measurement. Pixel resolution was converted to nanometers based on the magnification level. The fractional area that the calcium phosphate granules occupy per mitochondrion was calculated by multiplying the number of granules within a mitochondrion times the sum of all the granule areas divided by the area of the mitochondrion (N_granules_*A_granules_ / A_mito_). The calcium phosphate nanoclusters (n = 227) were determined by measuring the electron-dense regions located within the granules using ImageJ (NIH, Bethesda, MD, USA).

### Statistics

The Shapiro-Wilks test was used to confirm data normality. All data were analyzed and plotted using MATLAB 2019a (Mathworks, Inc., Natick, MA, USA). The data in figures 1 – 3 (n = 3 – 4) and stats presented for the calcium phosphate nanoclusters are presented as mean ± standard deviation. Mitochondrial images with calcium phosphate granules were only included for the histogram analysis (n value in the figures). An unpaired Student’s t test was used to compare the CsA treatment with the control group. An n-way ANOVA was run to determine significant effects between treatments at various calcium loads and different time-points. A p value < 0.05 was assumed to be statistically significant.

**Figure 1.**
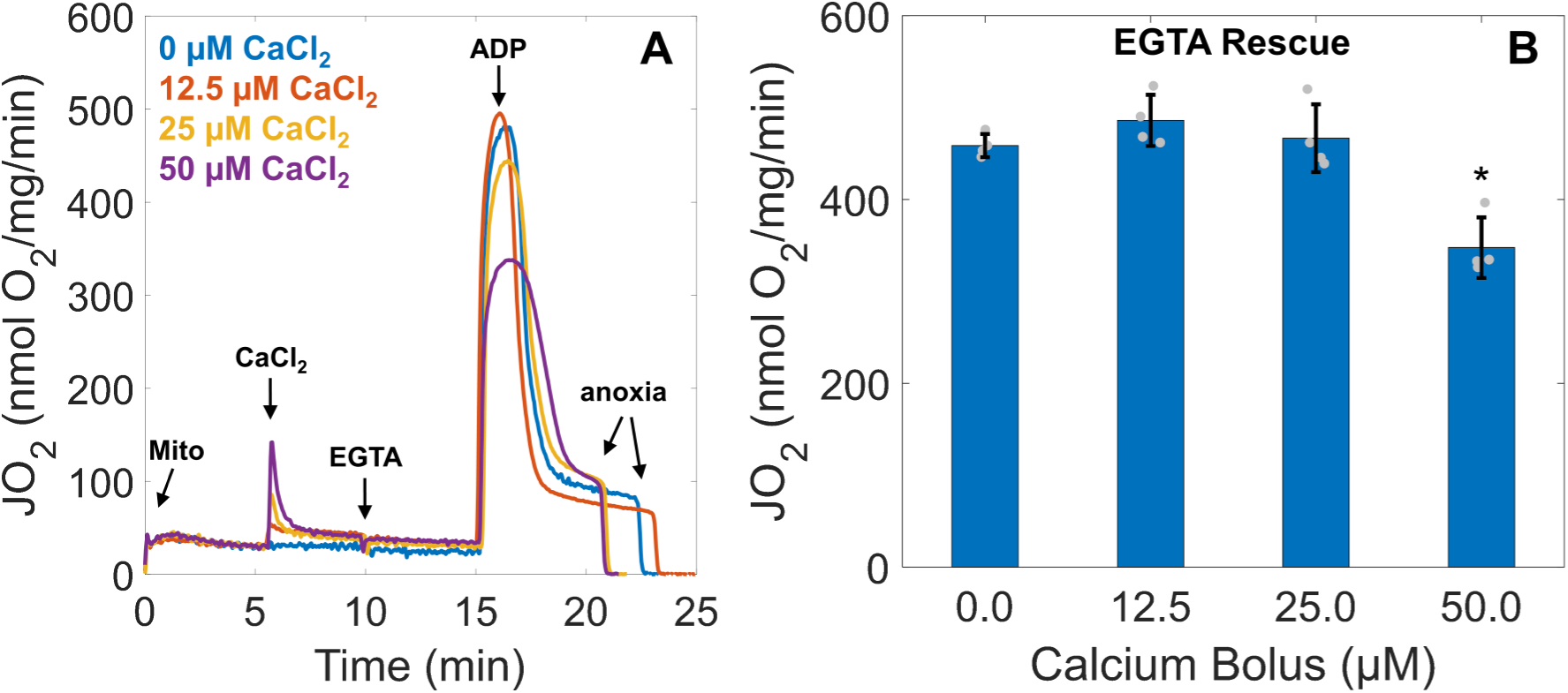
EGTA rescues mitochondrial function at low-to-moderate calcium loads but not high loads. A) Representative traces of ADP-stimulated respiration from calcium loaded mitochondria following the addition of EGTA. Mitochondria (0.1 mg/ml) were energized with 5 mM sodium pyruvate and 1 mM L-malate and exposed to various calcium boluses (0, 12.5, 25, and 50 µM). Five mins after calcium uptake, 1 mM EGTA was added to chelate all calcium in the system. Following an additional 5 mins, a bolus of 500 µM ADP was added to induce maximal ADP-stimulated respiration. B) ADP-stimulated mitochondrial respiration is recovered after EGTA addition for all but the 50 µM calcium bolus. Data are presented as mean ± standard deviation for a sample size of n = 4. Statistical comparisons are made with respect to 0 µM calcium. * represents a p value < 0.01.

### Reagents

All reagents were purchased from Sigma-Aldrich unless otherwise stated. Calcium Green™-5N hexapotassium salt was purchased from Thermo Fisher Scientific.

## Results and Discussion

### Respiratory inhibition by calcium overload is reversible in low-to-moderate calcium overload

While mitochondrial calcium concentrations lower than 100 nmol calcium/mg mitochondria supports ATP production (Bazil, Blomeyer, Pradhan, Camara, & Dash, 2013; Glancy et al., 2013; Wescott et al., 2019), levels above 500 nmol/mg mitochondria depress oxidative phosphorylation (Brustovetsky et al., 2003; Malyala, Zhang, Strubbe, & Bazil, 2019; Pandya, Nukala, & Sullivan, 2013; Territo, Mootha, French, & Balaban, 2000). In one of these studies, it was proposed that calcium phosphate precipitates form in the mitochondrial matrix at high calcium loads and reduce ATP production rates by either impeding metabolite transport and diffusion or destabilizing cristae, the functional units of mitochondria. However, the lasting effects of significant calcium accumulation were not explored in either of these studies. To test this, we monitored mitochondrial respiration rates following the addition of the calcium chelator EGTA under various calcium boluses in the range of 0 to 500 nmol/mg as shown in Figure 1.

The results in Figure 1 show that the inhibitory effect of calcium overload is reversible for all but high calcium loads. As expected, the respiratory rates – the amount of oxygen consumed per mitochondrial content at a given time – before calcium addition were equal across conditions. After a transient increase in respiration due to calcium uptake, respiration remains elevated because of the activation of calcium-sensitive matrix dehydrogenases and sodium/calcium cycling. When 1 mM EGTA was added to chelate buffer calcium, the ADP-driven respiratory rates were similar across all conditions except for the highest dose tested. These results suggest that when calcium overload exceeds a certain threshold, mitochondrial oxidative phosphorylation is irreversibly inhibited. This effect does not involve mitochondrial calpains (Malyala et al., 2019) and may involve some sort of structural change that lowers ATP production rates. Thus, the effect of calcium overload lies on a spectrum whereby higher levels of calcium result in detrimental changes in mitochondrial bioenergetic pathways.

### CsA preserves the mitochondrial function under high calcium loads

We then measured mitochondrial respiratory rates during excessive calcium overload by adding a 75 μM or 100 μM calcium bolus in the presence or absence of CsA, a known PTP inhibitor (Figure 2A and 2B). In agreement with results from Figure 1, increasing the extent of calcium overload impairs oxidative metabolism. However, the depressive effects of calcium on ADP-stimulated respiration is much more severe at these higher doses. The respiratory rate after ADP addition drops below 50 nmol O_2_/mg/min after the 75 μM CaCl_2_ bolus and drops below 20 nmol O_2_/mg/min for the 100 μM CaCl_2_ bolus. When CsA was present, this calcium-dependent inhibitory effect is partially mitigated with the rate reaching nearly 320 nmol O_2_/mg/min after the 75 μM bolus and 280 nmol O_2_/mg/min for the 100 μM bolus. Therefore, as others have found, CsA partially preserves mitochondrial function in the face of overwhelming calcium overload (Baines & Gutierrez-Aguilar, 2018; Bonora et al., 2016; De Marchi, Bonora, Giorgi, & Pinton, 2014; Fournier, Ducet, & Crevat, 1987; McGee & Baines, 2012). This effect is typically attributed to the ability of CsA to inhibit PTP opening; however, our structural data shown in the following sections suggest the existence of a novel protective effect of CsA.

**Figure 2.**
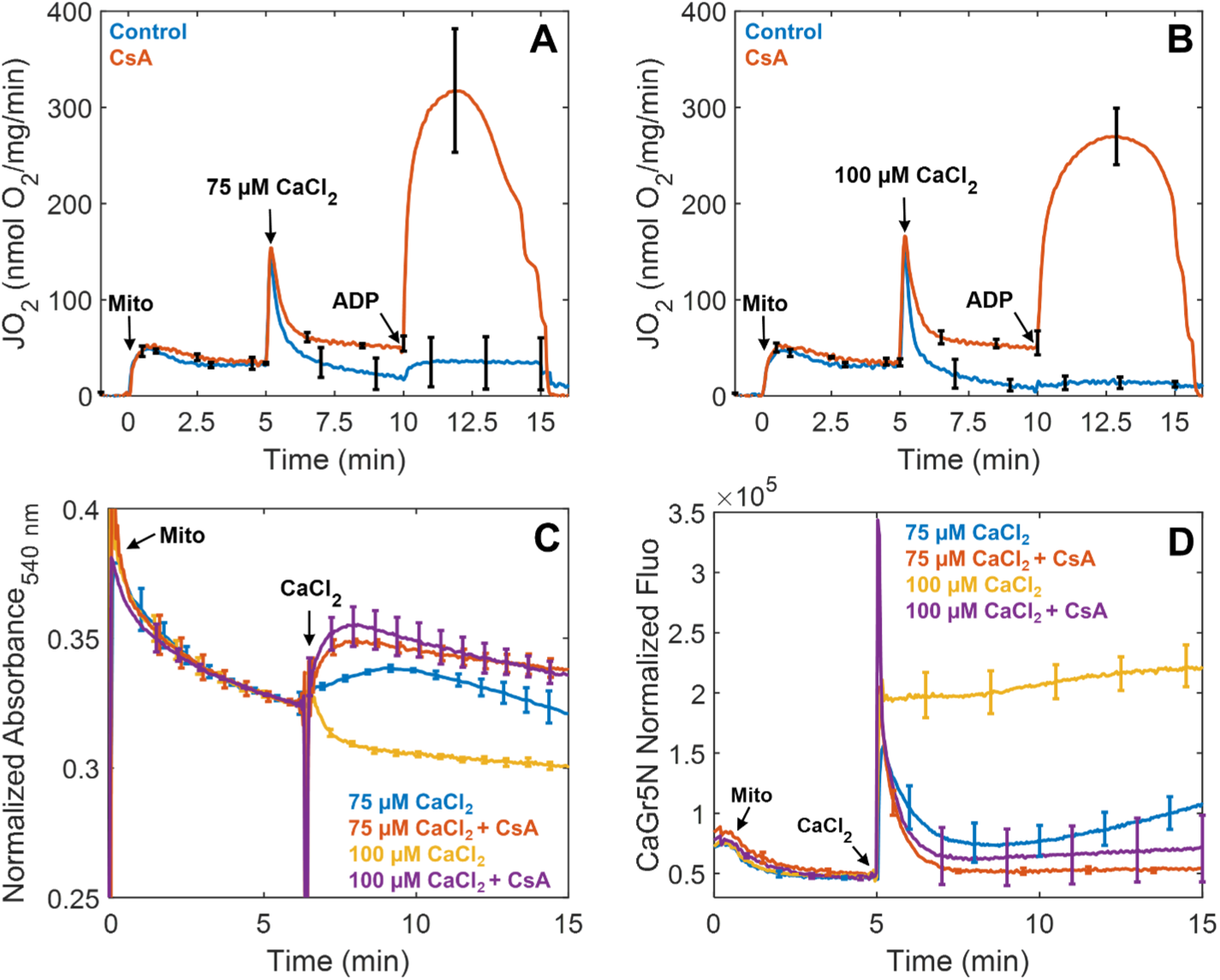
CsA preserves ADP-stimulated respiration, increase the absorbance, and enables robust calcium uptake in a calcium dependent manner. A and B) The addition of 1 µM CsA prevented a near-total collapse of ADP-stimulated respiration after a bolus of 75 or 100 µM calcium chloride. Mitochondria were energized as described in Figure 1. C) Mitochondrial swelling was monitored in parallel by quantifying absorbance at 540 nm. Large amplitude swelling was only observed in the 100 µM calcium bolus group when CsA was absent. D) Experimental conditions for calcium uptake were similar except that these experiments were performed in a cuvette open to atmosphere and tracked using the fluorescent probe CaGr5N (1 µM). In the absence of CsA, both calcium boluses were not completely taken up by the mitochondria, while in some instances, mitochondria can uptake calcium followed by release as shown after the addition of a 75 µM calcium chloride bolus. In contrast, CsA enables near-complete calcium uptake of either bolus. Data are presented as mean ± standard deviation for a sample size of n = 3 – 4.

In addition to the respirometry studies, mitochondrial absorbance data obtained in parallel (Figure 2C) shows that only the 100 µM calcium chloride bolus elicited large amplitude swelling, a classic indicator of mitochondrial permeability transition (Di Lisa, Menabò, Canton, Barile, & Bernardi, 2001). In contrast, the addition of a 75 µM calcium chloride bolus induced an increase in absorbance due to the formation of calcium phosphate granules scattering light at this wavelength (Chalmers & Nicholls, 2003). The gradual decrease in absorbance that follows is attributed to mitochondria fragmenting over time in response to the calcium insult. For both CsA-treated groups, the calcium-dependent increase in absorbance was sustained followed by a much slower decrease. However, the decrease in signal is not due to mitochondrial fragmentation but rather due to the inner membrane reorganization and matrix expansion (Beavis, Brannan, & Garlid, 1985; Garlid & Beavis, 1985). This phenomenon is discussed in latter sections of this report when classifying CsA-treated mitochondria. These results are similar to findings from a recent study that looked at the effects of the mitochondria-targeting peptide SS-31 on reducing infarct size of reperfused ischemic hearts (Brown et al., 2014). In that study, the changes in absorbance after CsA treatment following addition of calcium chloride are very similar to our results.

Our interpretation of the absorbance data is supported by our calcium uptake data shown in Figure 2D. These data also demonstrate the profound beneficial effects of CsA on mitochondrial calcium sequestration. When CsA was absent, mitochondria were not able to maintain calcium homeostasis and calcium was released into the buffer. For the 75 µM calcium chloride challenge, this release was gradual and suggests there is a snowball-like effect in which mitochondria with lower calcium tolerances release their calcium loads and force other mitochondria to take up even more calcium (Bernardi, 1992, 1999; Petronilli, Cola, Massari, Colonna, & Bernardi, 1993). This results in additional mitochondria losing their ability to retain calcium, and the process repeats until the entire mitochondrial population is compromised. In contrast, mitochondria were not able to effectively take up and store the 100 µM calcium bolus at all when CsA was absent. This level of calcium overload is sufficient to rapidly compromise the entire population in short order.

### Elucidating the effects on calcium overload on mitochondrial ultrastructure

To capture mitochondria undergoing MPT during calcium overload, we used the sampling scheme shown in Figure 3. These samples were drawn from a cuvette of isolated mitochondria at the indicated time points and subsequently vitrified in liquid ethane and imaged using cryo-EM. A total of 1345 cryo-EM images were analyzed and organized by sample time-point; before adding calcium (t_0_) and 1.5 mins (t_1_), 4 mins (t_2_), and 10 mins (t_3_) after adding a calcium bolus. We found that many mitochondria shared certain features at each time point and grouped them into 5 stages based on morphology and structure. Each stage represents the transition leading towards complete fragmentation and loss of function in the context of calcium phosphate granules abundance, growth, outer membrane rupture, cristae integrity, and inner membrane fragmentation (Table 1). An example of this stage classification is shown in Figure 4A. These panels represent the typical process induced by a 75 µM bolus of calcium in a population of isolated mitochondria.

**Table 1.** Mitochondrial Stages of Permeabilization Inferred from Structural Data

| Stage | Characterization |
| --- | --- |
| 1 | Rounded mitochondria with intact IMM/OMM with small or no granules present (n=33) |
| 2 | Large granules are present, initial localized OMM rupture, altered OMM morphology (n=59) |
| 3 | OMM partially or completely lost, beginning of IMM evisceration, granules begin to dissolve (n=48) |
| 4 | Significant IMM fragmentation, granules mostly dissolved (n=69) |
| 5 | Complete IMM fragmentation, only membrane fragments present, granules completely dissolved (n=65) |

OMM = outer mitochondrial membrane, IMM = inner mitochondrial membrane, n is the number of images collected with a mitochondrion in that stage

**Figure 3.**
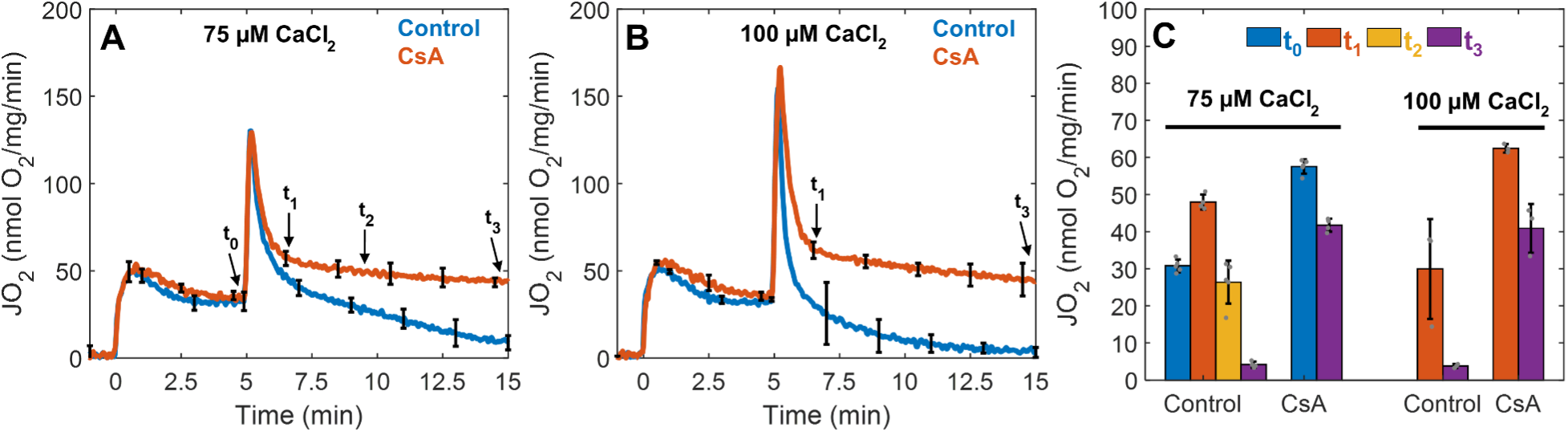
Cryo-EM sample collection protocol and time-points. A) For the 75 µM calcium chloride bolus, 5 µl of the mitochondrial suspension were collected and deposited on the Quantifoil holey-carbon grid at 5 mins just before calcium addition (t_0_), approximately 1.5 mins after calcium addition (t_1_), 4 mins after calcium addition (t_2_), and 10 mins after calcium addition (t_3_). B) For the 100 µM calcium chloride bolus, the mitochondrial suspension was sampled at t_1_ and t_3_. In all conditions, mitochondria (0.1 mg/ml) were energized with 5 mM sodium pyruvate with 1 mM L-malate. C) The effect of calcium in the presence or absence of CsA was quantified for each time-point. In the absence of CsA, mitochondrial respiration decreases dramatically as a function of time and the effect is exacerbated at greater calcium loads. In the presence of CsA, mitochondrial respiration was maintained. Data are presented as mean ± standard deviation for n = 3 – 5 biological replicates.

**Figure 4.**
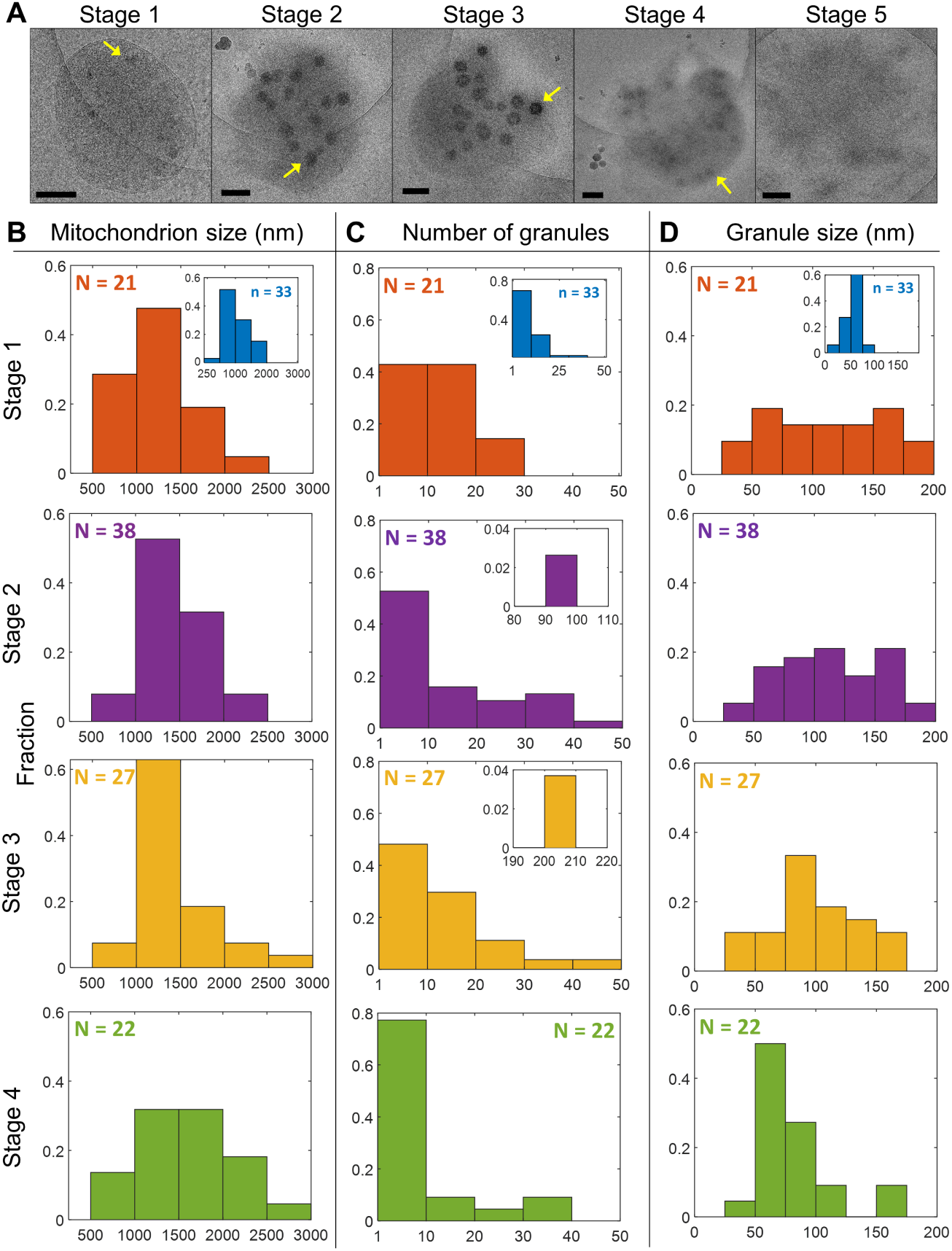
Calcium induces the formation of calcium phosphate granules, outer membrane rupture, and inner membrane evisceration. A) Representative images of mitochondria from stages 1 – 5 show that calcium induces inner membrane evisceration and outer membrane rupture. The lighter circle is the hole of the Quantifoil carbon grid. Mitochondria typically adhere to the carbon support film that is made hydrophilic after plasma treatment, so they are often either entirely on the carbon or half-on and half-off as shown in these images. The images from stage 2 through 4 contain ice contamination. These ice crystals appear as dark spots with a white fringe or halo outside and below or to the side of the mitochondria. These are easily distinguishable from calcium phosphate granules located in the mitochondrial matrix. Scale bars are 250 nm. The insets for the Stage 1 histograms (B-D) are histograms calculated from images collected before the addition of a 75 µM calcium bolus. The presence of these granules is due to ∼4 µM calcium contamination in the respiration buffer. The addition of a 75 µM calcium bolus leads to the formation of much larger and more abundant calcium phosphate granules of various sizes. B-D) The mitochondrial sizes and the calcium phosphate granules number and sizes per mitochondrion were further quantified. The mitochondrial size does not change significantly between stages. However, the granule abundance decreases by stage 4. Granules dissolving due to inner membrane fragmentation and loss of membrane potential cause this. The number of images analyzed for each stage is given by n. Note, no histograms for stage 5 are given as this stage consists of completely fragmented mitochondria with no calcium phosphate granules. The data from histograms were obtained from only mitochondria containing calcium phosphate granules. Arrows point to calcium phosphate granules.

Mitochondria in stage 1 have intact inner and outer membranes and are typically round (Figure 4A and S1). Cristae structures in this set of images are hard to distinguish; however, some are identifiable. Before the addition of a calcium bolus mitochondria are smaller and their inner and outer membranes remain intact as shown by the insets in Figure 4B – 4D. The number of calcium phosphate granules is relatively low with sizes averaging less than 100 nm in size. After the addition of 75 µM calcium chloride, mitochondria begin to fragment and lose bioenergetic competency. The beginning of this process is characterized by stage 2 (Figures 4 and S2). In this stage, regions of localized outer membrane rupture are observed and are always accompanied by the appearance of calcium phosphate granules. While the size of granules within a mitochondrion does not vary significantly, differences between mitochondria are common and noticeable (Figure S3). During the transition from stage 2 to stage 3, outer membrane definition is lost and the inner membrane is released. The inner membrane also begins to fragment in this stage. In some instances, calcium phosphate granules are still present indicating that the inner membrane is still energized. However, there are also images of this stage showing granules in the middle of dissolution (Figure S4), so this stage is when depolarization begins. Unexpectedly, the granules appear to dissolve from the inside out. In stage 4, the outer membrane is almost entirely gone, and inner membrane is extensively fragmented. Stage 5 is characterized by the complete fragmentation of the mitochondrial inner membrane and is the dominant stage at the 15 min time point. In this stage, mitochondria are deenergized and contain no calcium phosphate granules.

Interestingly, there were no large differences in mitochondrion sizes between the time points (ranging 500 – 3000 nm with mean values ∼1400 nm), but there were some clear differences in the size and number of granules (Figure 4B – 4D). As mitochondria transition from stage 1 to stage 3, the increase in absorbance shown in Figure 2C is caused by the increases in numbers and sizes of calcium phosphate granules. In fact, the number of calcium phosphate complexes per mitochondrion reaches a maximum by stage 3 and decreased in the following stage as shown in Figure 4C. The decreased in size and abundance by stage 4 is due to more complete mitochondrial permeabilization and fragmentation. Hence, for the first time to our knowledge, the MPT phenomenon now has direct visual confirmation of the processes proposed to occur. However, our results elucidate a mechanism that pinpoints cristae remodeling and inner membrane fragmentation as the key determinant of mitochondrial dysfunction as discussed further below.

### CsA preserves the inner membrane, promotes the formation of granules of greater size, and increases the abundance

Next, we repeated the calcium overload imaging experiments in the presence of CsA to understand how mitochondrial respiration and calcium handling were preserved from an ultrastructural perspective (Figure 5 and S5). Similar to control mitochondria, CsA-treated samples were grouped into 4 classes based on morphology (Table 2). However, the classes are not related to a sequence of events like the stages, rather they are descriptive. Many of the images showed normal looking mitochondria with well-defined inner and outer membranes. These are class 1 mitochondria. Some of these mitochondria contained granules caused by uptake of low levels of contaminant calcium. In addition, some mitochondria had a condensed inner membrane that was sometimes localized to one side of the mitochondrion. These electron-dense regions are presumably area of high cristae density. Interesting, some images showed mitochondria with the outer membrane ruptured with the inner membrane partially or more completely ejected from the mitochondrion. These mitochondria are classified as class 2 mitochondria. In other images, mitochondria were clustered together and are defined as class 3 mitochondria. Lastly, after the calcium treatment, images revealed mitochondria with no outer membrane, large calcium phosphate granules, and the inner membrane spread across the carbon grid. These mitochondria are classified as class 4 mitochondria. Because of morphological changes induced by CsA and calcium addition, the sizes of these mitochondria are larger than mitochondria in the other classes. In addition, mitochondria in this class had granules of heterogeneous sizes between but rarely within a single mitochondrion (Figure S6). Despite these radical changes in ultrastructure, the mitochondria remain functionally competent as shown in Figure 2. The best explanation for this observation is that the cristae junctions and inner membrane integrity is preserved by the CsA treatment.

**Table 2.** Mitochondrial Classes of Inferred After CsA Treatment

| Class | Characterization |
| --- | --- |
| 1 | Normal looking mitochondria with defined OMM, condensed IMM, and with or without granules (n=215) |
| 2 | Inner mitochondrial membrane ejection (n=48) |
| 3 | Clustered mitochondria (n=81) |
| 4 | IMM spread out but intact with granules present (n=219) |

OMM = outer mitochondrial membrane, IMM = inner mitochondrial membrane, n is the number of images collected with a mitochondrion in that stage

**Figure 5.**
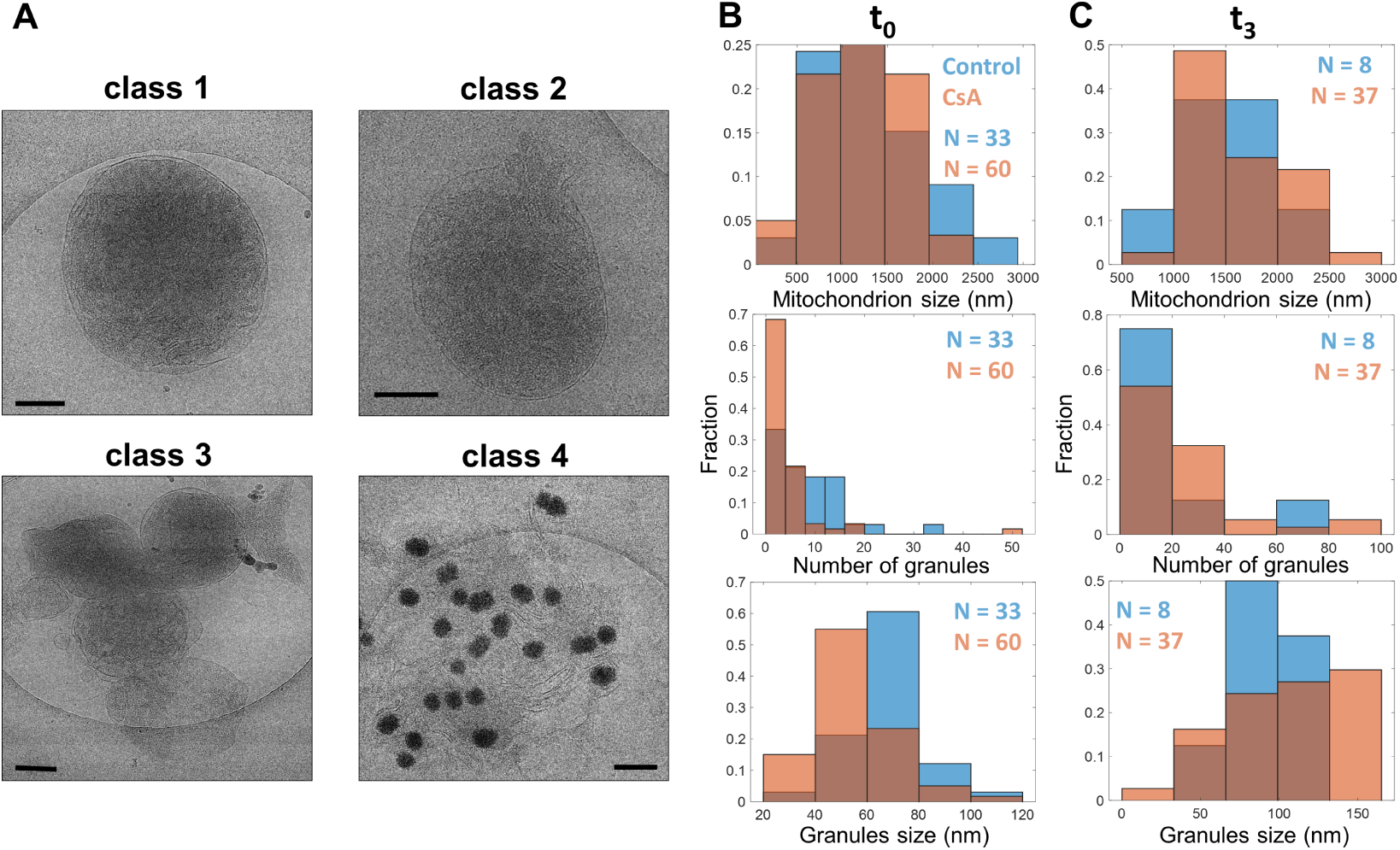
CsA disrupts the outer membrane morphology, causes release of the inner membrane, tends to form mitochondrial clusters, and enhances the number and size of calcium phosphate granules per mitochondrion. Representative images before the addition of a 75 µM calcium bolus (t_0_) in the presence of 1 µM CsA. Mitochondria were energized with 5 mM sodium pyruvate and 1 mM L-malate. A) CsA induced morphological changes to mitochondria that can be grouped into 4 classes as described in Table 2. B) The mitochondrial size, calcium phosphate granules size and number per mitochondrion were quantified for each time-point (t_0_ – t_3_) before and after the addition of a 75 µM calcium bolus in the presence or absence of CsA. B) There are no differences in the mitochondrial size of control to CsA-treated mitochondria before the addition of calcium. C) After the calcium addition, the number and size of the granules increased in CsA-treated mitochondria were much larger than in the control mitochondria. Scale bars are 250 nm. n represents the number of images analyzed by the time point for control and CsA treated conditions.

After calcium addition, the abundance and size of granules per mitochondrion and the mitochondrion size increased in the presence of CsA compared to the control group (Figure 5). Before the addition of calcium, the average control mitochondrion size was 1320 ± 550 nm, the average granule size of 68 ± 14 nm, and the average number of granules per mitochondrion was 9.7 ± 3.1. Whereas the average CsA-treated mitochondrion size was 1180 ± 400 nm, the average granule size was 55 ± 16 nm, and the average number of granules per mitochondrion was 5.1 ± 2.3. These results show that CsA does not influence any of these parameters before the large calcium bolus was administered. However, after calcium addition, there are noticeable differences between control and CsA-treated mitochondria. The mitochondrial size for control mitochondria average 1470 ± 530 nm with a granule size of 90 ± 22 nm and abundance of 18.0 ± 4.3 per mitochondrion. Whereas CsA-treated mitochondria size average was 1630± 400 nm with a granule size of 102 ± 36 nm and abundance of 26.0 ± 5.1 per mitochondrion. Of note, only a small number of control mitochondria survived by the last time point (Figure 5C).

### Mitochondrial membrane fragmentation occurs more rapidly at greater calcium loads but is mitigated by CsA

Seeking to understand the observed large amplitude swelling for the control group after the addition of a 100 µM calcium chloride bolus, images of control mitochondria were collected after a 100 µM calcium bolus addition. Most of the images displayed outer membrane rupture at multiple regions suggesting a rapid expansion of the inner membrane compared to the 75 µM calcium bolus (Figures 6, S7 and S8). Thus, at this high of a calcium bolus, the morphological changes were caused by what appears to be *bona-fide* permeability transition pore opening. As expected, CsA prevented this rapid expansion and led to the formation of numerous and large calcium phosphate granules. Without CsA, the size and abundance of the granules were noticeably decreased (Figure 6B). While there were no differences in mitochondrion sizes between treatments shortly after calcium addition (1320 ± 370 nm vs 1490 ± 380), the average control mitochondrial size decreased to 1160 ± 440 nm in the last time point (Figure 6C). In contrast, the average CsA-treated mitochondrion size increased to 1710 ± 440 nm. The average number of granules in control mitochondria as a function of time was reduced from 9.6 ± 3.1 to 6.6 ± 2.6. The average size of these granules marginally increased from 84 ± 32 to 90 ± 37 nm. The average number of granules in CsA-treated mitochondria increased from 46.1 ± 6.8 to 107 ± 10 with average sizes increasing from 121 ± 21 to 132 ± 28 nm. These values are greater compared to the values measured after a 75 µM calcium bolus was given. This is consistent with CsA increasing calcium accumulation and preserving mitochondrial function even at these high calcium loads. However, the oxygen consumption rate after the 100 µM calcium bolus was significantly lowered compared to the 75 µM calcium bolus (Figure 2A and 2B). Hence, we conclude that calcium induces irreversible effects on mitochondrial function and CsA, although not entirely protective, delays complete loss of function and allows more calcium uptake.

**Figure 6.**
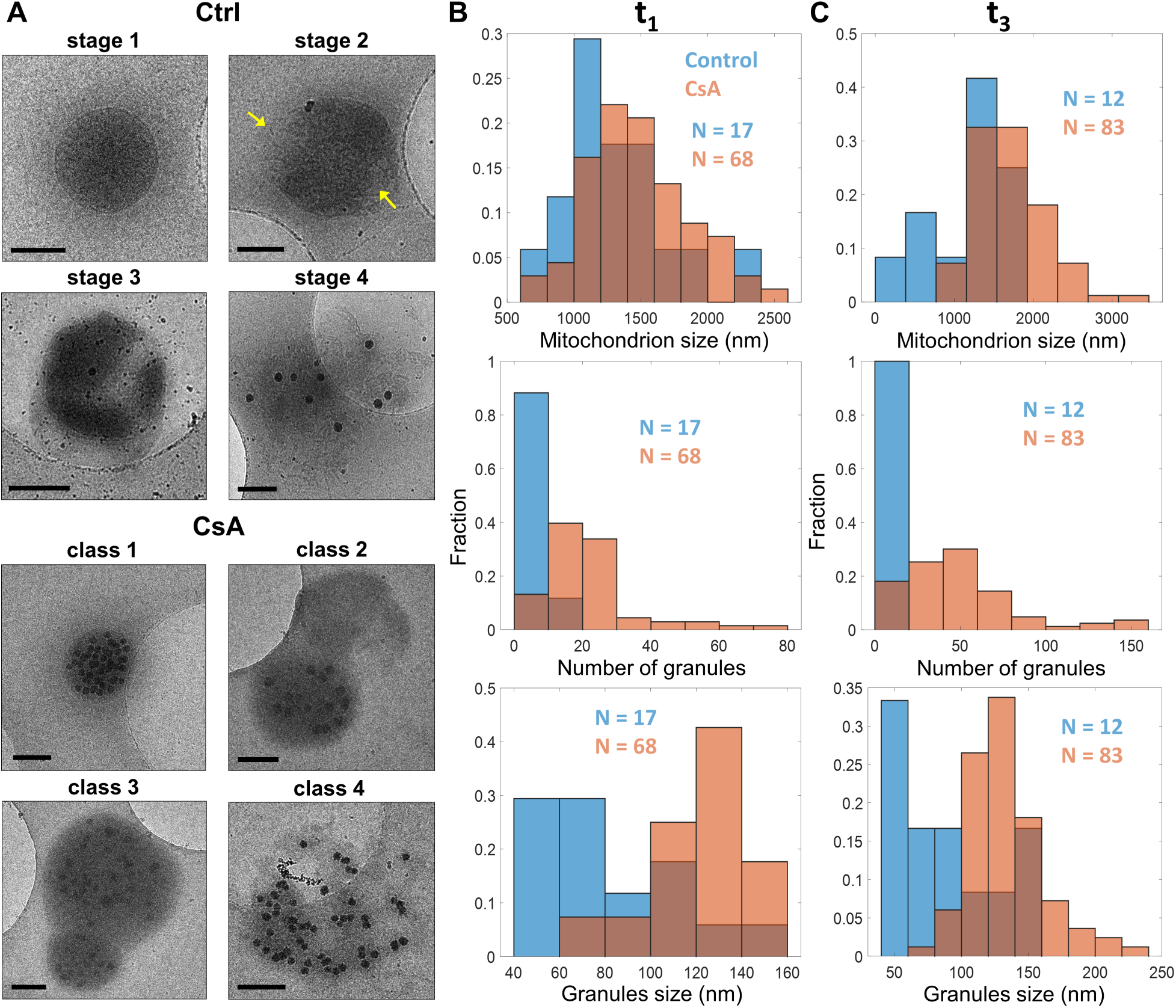
In extreme calcium overload conditions, CsA preserves inner mitochondrial membrane intactness. Representative images before the addition of a 100 µM calcium bolus in the presence or absence of 1 µM CsA. A) The addition of 100 µM calcium led to the formation of granules of various sizes varying between but not within mitochondrion. This large bolus of calcium induces membrane rupture in control mitochondria in multiple location as shown by the arrows. However, mitochondria treated with CsA were protected. B) There is no differences in the average mitochondrial size just after calcium addition between control and CsA-treated mitochondria, but the CsA-treated mitochondria had larger and more abundant granules. C) At 10 mins after the addition of calcium, all three measures (mitochondrial size, granule size, granule number) become larger in the CsA-treated mitochondria. In contrast, all three measures decreased in control mitochondria due to membrane fragmentation and evisceration. The number of images analyzed by the time point for control and CsA treated conditions is given by n. Scale bars are 250 nm.

### Calcium phosphate granules are composed of smaller structural units

Calcium phosphate complexes are considered the main component of the mitochondrial calcium sequestration system (Malyala et al., 2019; Nicholls & Chalmers, 2004; Wolf et al., 2017). Pioneering studies by Posner and others suggested that amorphous calcium phosphate consists of many smaller spherical elements with a chemical composition of Ca_9_(PO_4_)_6_ (Eliaz & Metoki, 2017; Jiang, Pan, Chen, Xu, & Tang, 2015; Mancardi, 2017; Onuma, 1998). These elementary calcium phosphate units were named as Posner clusters with a diameter ranging from 0.7 to 1.0 nm (Onuma, 1998). In the present study, we lack the image resolution to resolve individual Posner clusters. However, our data show that the calcium phosphate granules are composed of highly electron-dense regions that resemble Posner clusters stacked together forming a higher order granule structure as shown in Figure 7. Our data also reveal that these structures are independent of CsA treatment or sampling time with the only major difference between clusters being their size. Assuming that these higher order granule structures are packed in a body-centered cubic lattice, the volume fraction of solid calcium phosphate is approximately 70%. Computing the total volume fraction of these granules by multiplying the volume of individual granules and their numbers per estimated mitochondrial volume (based on measured diameters and approximate thickness of 300 nm) gives an average volume fraction of 0.025. The thickness is a constraint applied after thinning the water layer before vitrification. Then by assuming a Posner cluster volume of 0.32 nm^3^, the average calcium load per mitochondrion is 796 nmol/mg. To convert from mitochondrial volume to mitochondrial mass, we used the estimate of 6×10^9^ mitochondria/mg reported in Beavis and Garlid (Beavis et al., 1985). This value, 796 nmol/mg, is strikingly similar to the expected value of 750 nmol/mg calculated from the calcium uptake data shown in Figure 2D. Thus, these images yield expected values of calcium uptake and show the overwhelming majority of calcium taken up by mitochondria is stored in these calcium phosphate granules.

**Figure 7.**
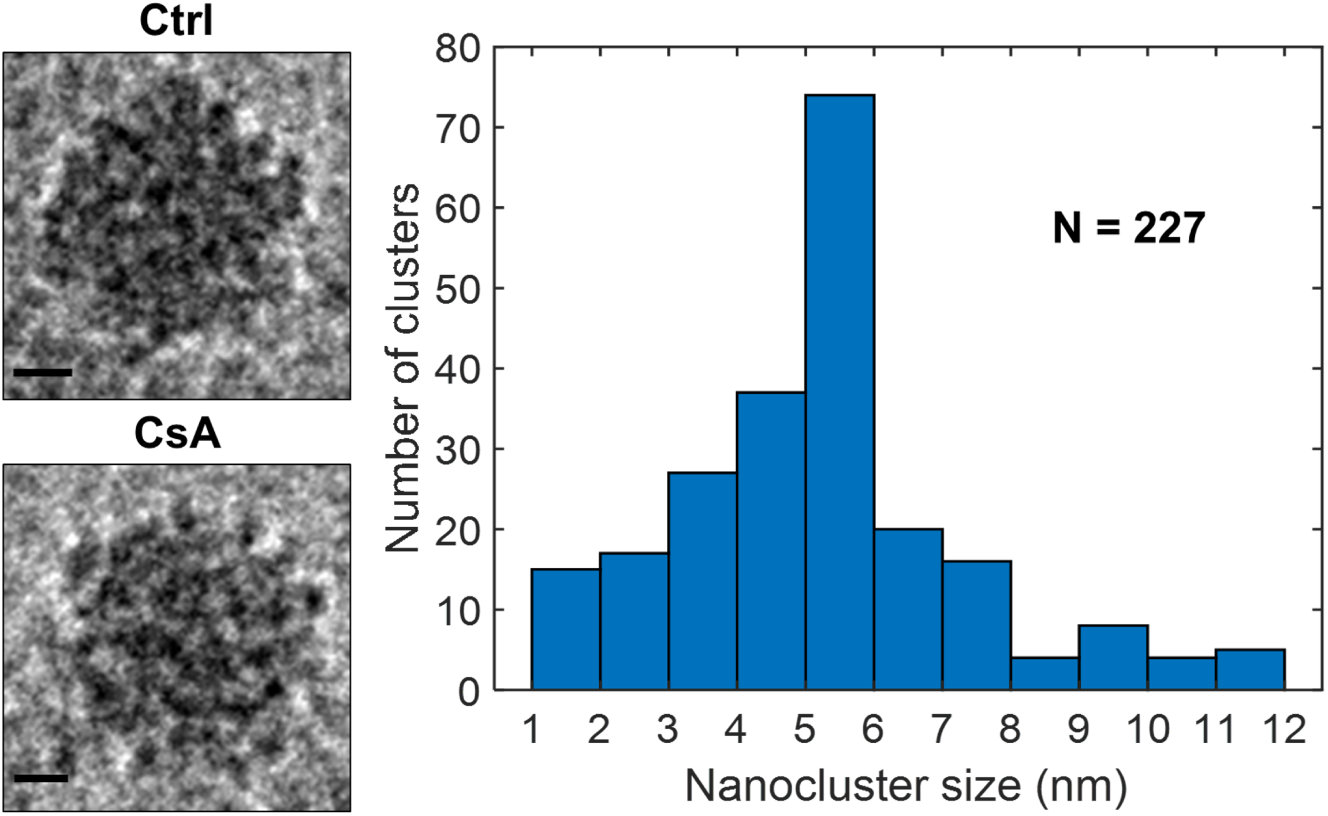
Calcium phosphate granule structure. Left) Representative images of calcium phosphate granules for each calcium condition in the presence or absence of CsA shows near identical structure. Right) Each granule consists of many individual electron-dense calcium phosphate nanoclusters with a diameter of 5.3 ± 2.1 nm. Scale bars are 25 nm.

While the vast majority of the isolated mitochondria contained calcium phosphate granules following the calcium addition, a few did not. There are two possible explanations for this phenomenon; either this group of mitochondria is 1) de-energized preventing calcium uptake or 2) they lack mitochondrial calcium uniporters (MCU). Based on the following statistical arguments, the latter is a more likely explanation. Assuming there are 40 MCUs per mitochondrion (Wescott et al., 2019) with an estimated standard deviation of 20, the probability of randomly selecting a mitochondrion without an MCU channel is 2.3 %. This corresponds to 11 mitochondria in our total set of 502 images. In agreement with this estimation, our data show that 17 mitochondria do not possess granules after either calcium bolus was given which corresponds to 3.4 % of the number of mitochondria imaged. This percentage is independent of treatment with 3.5 % of control mitochondria and 3.6 % of CsA treated mitochondria without any calcium phosphate granules. These results also match the respirometry data given in Figure 2 whereby even after a large bolus of calcium chloride, some mitochondria are bioenergetically competent and synthesize ATP after the ADP bolus. Assuming that the boluses of calcium were sufficient to elicit MPT in the mitochondria with an MCU channel, the measured ADP-stimulated respiratory rate increase must be due to mitochondrial lacking an MCU. In line with this observation, the maximum ADP-stimulated respiratory rate for each calcium treatment relative to the maximum rate without calcium (as shown in Figure 1), is 10.9 % ± 5.2 and 4.3 % ± 1.1 for the 75 µM and 100 µM calcium bolus, respectively. These values are strikingly close to the value estimated from the imaging data.

### A new sequence of events that leads to mitochondrial dysfunction

The current leading hypothesis of calcium-induced mitochondrial dysfunction involves the peptidyl-prolyl *cis-trans* isomerase, cyclophilin D (CypD), interacting with as yet to be identified inner membrane proteins to form the permeability transition pore (Baines & Gutierrez-Aguilar, 2018; Bernardi, Di Lisa, Fogolari, & Lippe, 2015; Crompton et al., 1988; Kim et al., 2014; Porter & Beutner, 2018). When open, the pore results in sustained membrane depolarization, large amplitude swelling, and calcium release (Elmore, Qian, Grissom, & Lemasters, 2001; D. R. Hunter & Haworth, 1979), and loss of mitochondrial respiratory control. However, CsA can bind to CypD and sequester it so that its interaction with its target is prevented (De Marchi et al., 2014). However, CsA is not fully protective. It is believed that it only increases the calcium threshold required to open the pore. This idea is based on studies that show CsA increases the calcium retention capacity by nearly 3-fold (Chalmers & Nicholls, 2003). While an attractive hypothesis, this model has problems that are easier to explain using a different mechanism. As an alternatisve, we propose a novel mechanism of action whereby CsA enables robust calcium accumulation in the context of promoting calcium uptake and calcium phosphate granule formation. This mechanism involves the interaction between putative CsA-regulated proteins, and cristae structural proteins to preserve the inner membrane intactness. While the calcium phosphate granules may induces changes in morphology by mechanically disrupting membranes, it is plausible that free calcium interacts with proteins regulating inner membrane and cristae maintenance (namely the optic atrophic factor 1 and the mitochondrial contact site and cristae organizing system; known as OPA1 and MICOS) or additional regulators of this system. For instance, the stress-sensing overlapping activity with *m*-AAA protease 1 (OMA1) is a zinc metallopeptidase found in the inner mitochondrial membrane regulating mitochondrial dynamics through OPA1 processing (Consolato, Maltecca, Tulli, Sambri, & Casari, 2018; Opalinska & Janska, 2018; Rainbolt, Lebeau, Puchades, & Wiseman, 2016). OMA1 is activated under stress conditions including membrane potential dissipation, decreased ATP levels, and oxidative stress, among other insults (Consolato et al., 2018). Upon activation, OMA1 cleaves the long isoform inducing cristae remodeling and cyt. c release (Arnoult, Grodet, Lee, Estaquier, & Blackstone, 2005; Frezza et al., 2006; Glytsou et al., 2016; Rainbolt et al., 2016; Scorrano et al., 2002). This mechanism can explain the morphological and functional changes included by calcium overload that we observed in our cryo-EM images and bioenergetics data.

Indeed, we demonstrated that calcium overload impairs mitochondrial ATP production at greater calcium loads and depleting mitochondria of calcium did not fully restore function - indicating an irreversible component. These data revealed an underappreciated energetic consequence of calcium overload on the mitochondrial function that supports a direct role of the mitochondrial calcium buffering system. In cardiac tissue, the steady-state cycling of calcium across plasma membranes maintains cytosolic calcium levels at ∼100 nM during diastole; however, the peak calcium concentration during systole can rise to the low micromolar range (Balaban, 2002; Bazil et al., 2013; Boyman et al., 2014). Whether the mitochondria can respond to these transient changes to meet metabolic demand is a subject of debate (reviewed in (Balaban, 2002)) that revolves around the mitochondrial calcium uniporter (MCU) being unable to approach maximum flux rates in the transient rise of cytosolic calcium due to its low affinity for calcium (Marchi & Pinton, 2014). Alternative hypotheses regarding calcium microdomains have been proposed in an attempt to argue in favor of significant mitochondrial calcium uptake during systole (Csordás, 2001; De La Fuente et al., 2018; De la Fuente & Sheu, 2019; Garcia-Perez, Hajnoczky, & Csordas, 2008; Hom & Sheu, 2009); however, direct imaging studies do not support this (Boyman et al., 2014; Lukyanenko, Chikando, & Lederer, 2009). A recent study by Wescott et. al found that physiological cytosolic calcium transients causes a gradual, step-wise increase in matrix calcium concentration per beat rather than large transient peaks (Wescott et al., 2019). They also showed that at high pacing rates, the matrix calcium concentration did not change any further. Indeed, further studies are required to determine whether these results are due to equal influx and efflux of calcium per cycle or due to calcium buffering. At this point, we believe the calcium buffering in the form of calcium phosphate granules system becomes relevant. In a separate study, calcium phosphate granule deposits were observed in the matrix near cristae junctions in a variety of different eukaryotic cells under physiological conditions (Wolf et al., 2017). Given the relevance of calcium in bioenergetics, the presence of these calcium deposits may exert some degree of control over mitochondrial signaling and metabolism.

In Figure 8, we present a model that accounts for various characteristics of membrane fragmentation before the MPT onset. This model integrates findings from our cryo-EM analysis with mitochondrial function and recapitulates the effects of calcium on the mitochondrial structure. Based on our findings, we believe that changes in mitochondrial ultrastructure can explain the loss of mitochondrial function in calcium overload as well as the protective effects of CsA. Our results suggest that mitochondrial outer membrane rupture and inner membrane fragmentation are *caused* by calcium overload whereas the formation of granules are a *consequence* of calcium uptake and accumulation. In the present study, the detrimental effects of calcium overload on mitochondrial function are mitigated when CsA is present. Regardless of the calcium bolus, the number and size of granules in CsA-treated mitochondria increased, suggesting that CsA increases the mitochondrial calcium buffering capacity, thus explaining why CsA allows robust calcium uptake and increases the threshold for permeability transition pore activation (Baines & Gutierrez-Aguilar, 2018; Bonora et al., 2013; De Marchi et al., 2014; Fournier et al., 1987).

**Figure 8.**
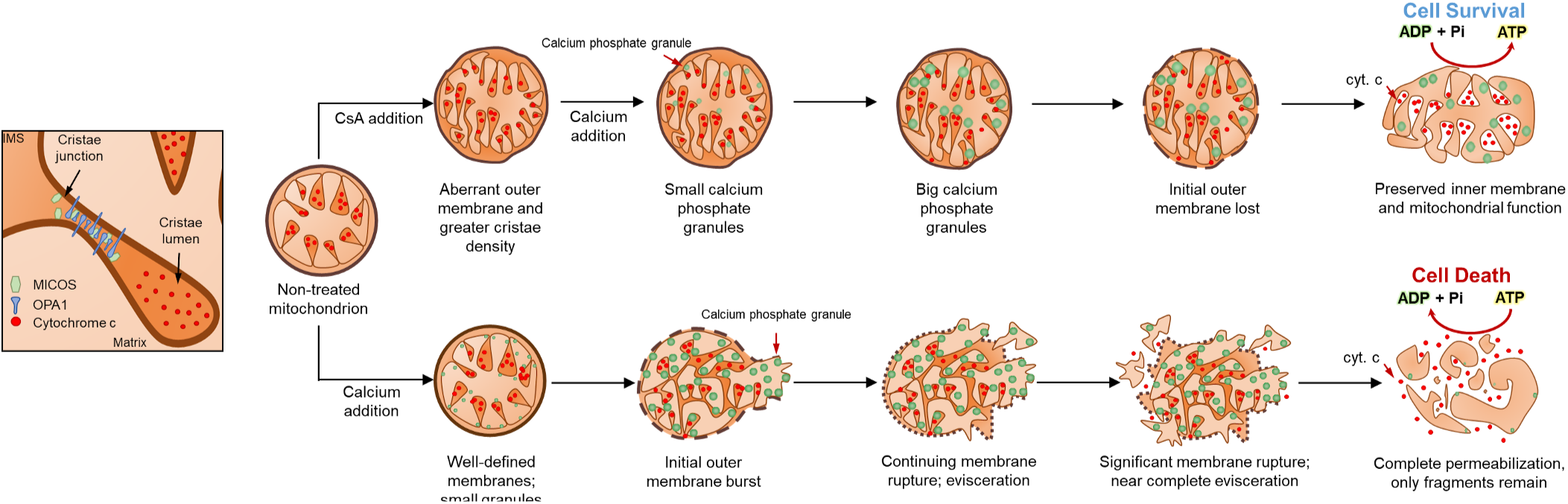
Schematic representation of calcium overload leading to mitochondrial fragmentation and permeabilization. In energized mitochondria, the mitochondrial membrane potential creates the driving force for calcium to accumulate in the mitochondrial matrix. The accumulation and growth of these complexes induces mitochondrial swelling that leads to outer membrane rupture, inner membrane fragmentation and cyt. c release. This causes membrane potential dissipation and induces the calcium-phosphate complex disassembly. CsA, however, alters the membrane morphologies and allows for robust calcium uptake after the addition of a calcium bolus. The inner mitochondrial membrane remains tethered and the cristae junctions intact. This avoids cyt. c remodeling and preserves the bioenergetic status despite the calcium effects on mitochondrial respiration.

To interpret these results, we sought confirmation of our findings from work by others. A study by Pinton’s group (Bonora et al., 2016) studied the effect of calcium overload on mitochondria in HeLa cells. Exposing HeLa cells to the ionophore ionomycin resulted in mitochondrial network fragmentation. However, in the presence of CsA, the mitochondrial network condensed and maintained its integrity after ionomycin treatment. In addition, a study looking at mitochondrial swelling using light transmittance in a single mitochondrion showed that calcium induces mitochondrial swelling in a concentration-dependent fashion (Shibata, Yoneda, Morikawa, & Ohta, 2019). CsA decreased this effect in a calcium-dependent manner, which led the authors to conclude that either CsA induces mitochondria shrinkage or calcium accumulation induces light scattering. We show that CsA increases the absorbance in a calcium-dependent manner and induces changes in mitochondria ultrastructure including condensed inner membranes and loss of outer membrane that often led to inner membrane unraveling. Therefore, our results are consistent with these studies but quantitatively describe the ultrastructural changes associated with calcium overload and how these changes are linked to mitochondrial function.

A major challenge in this study is the lack of cristae structural definition in our set of images. As dynamic structures, cristae are the functional units of mitochondria that lock cyt. c in the cristae lumen and provide sufficient membrane surface area to sustain oxidative phosphorylation at high rates (Mannella, 2006; Scorrano et al., 2002). Under certain conditions when the cristae junctional width is enlarged, cyt. c escapes the lumen and causes loss of mitochondria function and cell death (Hessenberger et al., 2017; Ikon & Ryan, 2017). While the expected outcomes during calcium overload were addressed, intricate details of the cristae structure including junction width, length, density, and shape must be incorporated to better understand the implications of cristae remodeling as key mediators in mitochondrial function. Earlier studies looking at calcium phosphate granule composition relied on staining, fixing or dehydrating samples, which introduce artifacts and make them less reliable (Brighton & Hunt, 1976; Carafoli, Rossi, & Lehninger, 1964; Greenawalt, Rossi, & Lehninger, 1964; Kristian, Weatherby, Bates, & Fiskum, 2002; Matthews & Martin, 1971). More recently, changes in mitochondrial structure were analyzed using high-pressure techniques and freeze-substitution to minimize sample structural distortion resulting from fixation or dehydration (Kristian, Pivovarova, Fiskum, & Andrews, 2007). However, structural details such as granule space distribution and structure are not as well defined with this method relative to the latest advanced cryo-EM techniques. Hence, visualization of the mitochondria in 3D by cryo-electron tomography (cryo-ET) would be an avenue for future studies to address. Nonetheless, our finding that CsA preserves the inner membrane integrity suggests that cristae remodeling and cyt. c release from the cristae lumen is likely avoided. This poses a new approach by which therapies targeting cristae remodeling can be identified to prevent pathological mitochondrial dysfunction leading to tissue injury.

## Conclusion

Mitochondrial calcium overload causes mitochondrial respiratory dysfunction through the well-established mitochondrial permeability transition phenomenon. However, the precise and quantitative causal mechanisms are only recently coming into focus. In this study, we have shown that calcium-induced inhibition of ADP-stimulated respiration is reversible for low-to-moderate calcium loads (0 – 250 nmol/mg) but only partially reversible for higher calcium loads (> 500 nmol/mg). Accumulating large quantities of calcium leads to the formation of calcium phosphate precipitates, outer membrane rupture, and eventually inner membrane fragmentation and evisceration. This results in loss of respiratory control and poor ATP synthesis rates. However, in the presence of CsA, the mitochondrial outer membrane is lost, but the ability of mitochondria to sequester significant amounts of calcium is retained with more abundant and larger calcium phosphate granules that persist. Here we conclude from our functional and structural data that CsA preserves the inner membrane integrity and mitochondrial function by preventing detrimental cristae remodeling associated with calcium overload. These findings mechanistically link mitochondrial function to ultrastructure in the context of calcium overload and brings a new understanding of the calcium sequestration system in relation to energy metabolism, structure, and potential targets to prevent mitochondrial dysfunction.

## Acknowledgments

The authors thank Dr. Alexander Makhov for technical support with cryo-electron microscopy. This work was supported in part by the Office of the Director, National Institutes of Health, under Award Number S10 OD019995 (JFC). The content is solely the responsibility of the authors and does not necessarily represent the official views of the National Institutes of Health. The AAAS Marion Mason Milligan award provided funding for this project for Women in the Chemical Sciences, the JK Billman, Jr., MD Endowed Research Professorhip, and NIH R01 GM110185 (KNP).

## Declaration of Interests

The authors declare no competing interests.

## Notes

#### Summary of Updates

This revised version, version 2, contains a correction in page 13, line 430. Lederers group was changed to Wescott et al.

