## Supplemental for "The Mitochondrial Permeability Transition Phenomenon Elucidated by Cryo-EM Reveals the Genuine Impact of Calcium Overload on Mitochondrial Structure and Function"

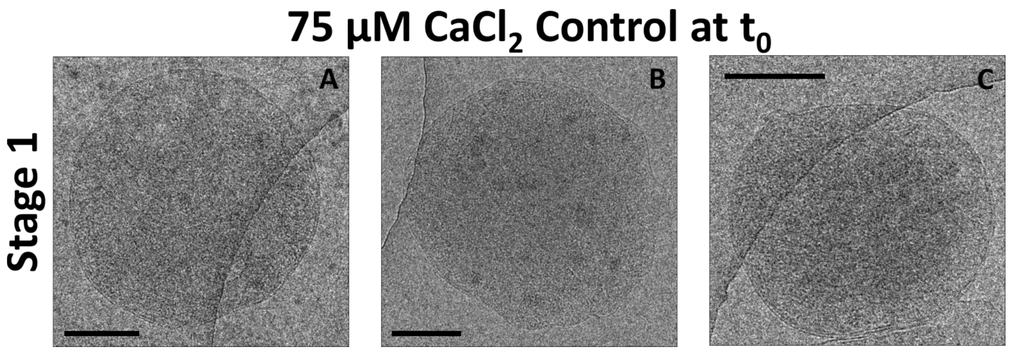


**Figure S1. Mitochondrial membrane morphology in the absence of calcium overload.** Representative images before the addition of calcium in the absence of CsA. A-C) Mitochondria are round with the outer and inner membrane clearly visible. In some instances, small calcium phosphate granules are observed. Granules form after mitochondria take up contaminate calcium. Samples were collected at $t_0$ as described in Figure 3. Scale bars are 250 nm.

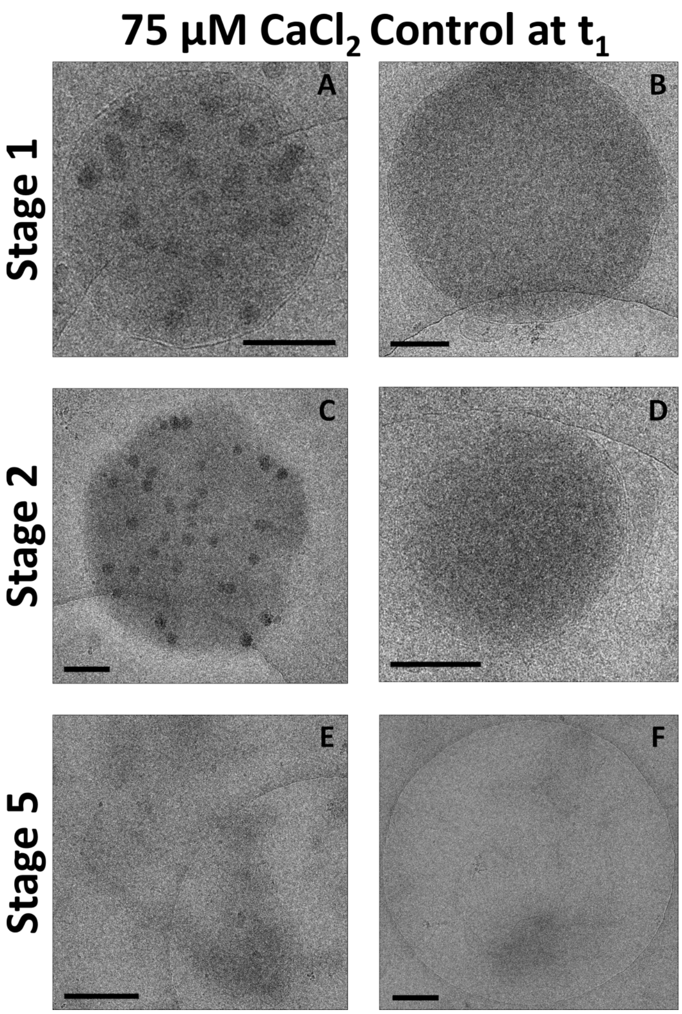


**Figure S2. Calcium uptake leads to the formation of large calcium phosphate granules, fragmentation the inner mitochondrial membrane, and rupture of the outer membrane** Representative images of 1.5 mins ($t_1$) after the addition of a 75 µM calcium chloride bolus. A and B) Mitochondria with preserved inner and outer membranes observed. C and D) The beginning of mitochondrial fragmentation caused by calcium overload. E and F) Absence of outer membrane with fragmented inner membranes. Scale bars are 250 nm.

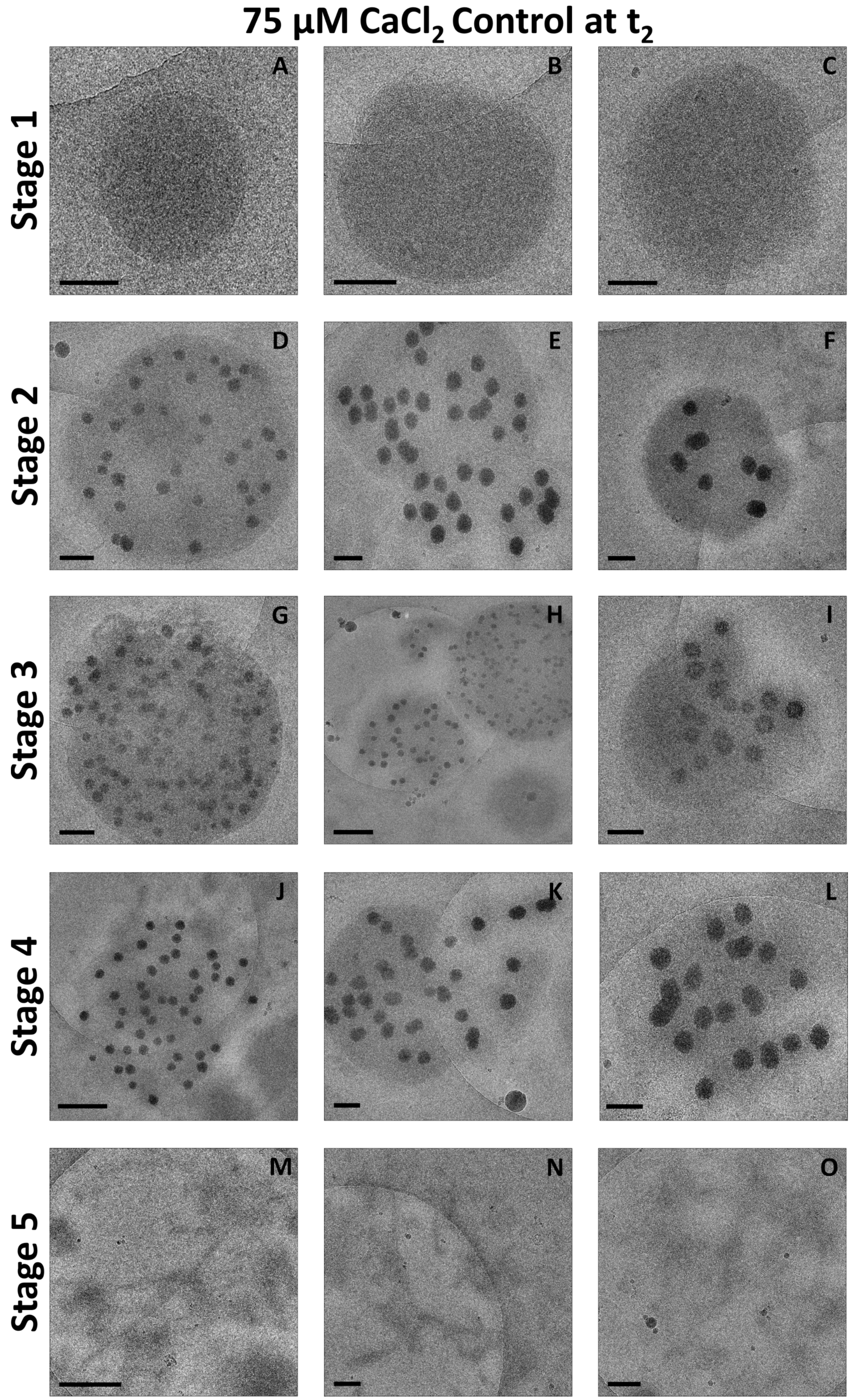


**Figure S3. Calcium accumulation induces localized ruptures of the outer membrane and complete mitochondrial fragmentation.** Representative images of 4 mins ($t_2$) after the addition of a 75 μM calcium chloride bolus. A-D) Defined inner and outer membranes with granules present or absent. D-F) Loss of outer membrane integrity. G-I) Loss of the outer membrane begins initiating inner membrane fragmentation. J-L) Complete loss of outer membrane with preserved granules. M-O) Complete loss of inner and outer membranes. In this stage, the inner membrane is severely compromised and unable to maintain a membrane potential. Scale bars are 250 nm.

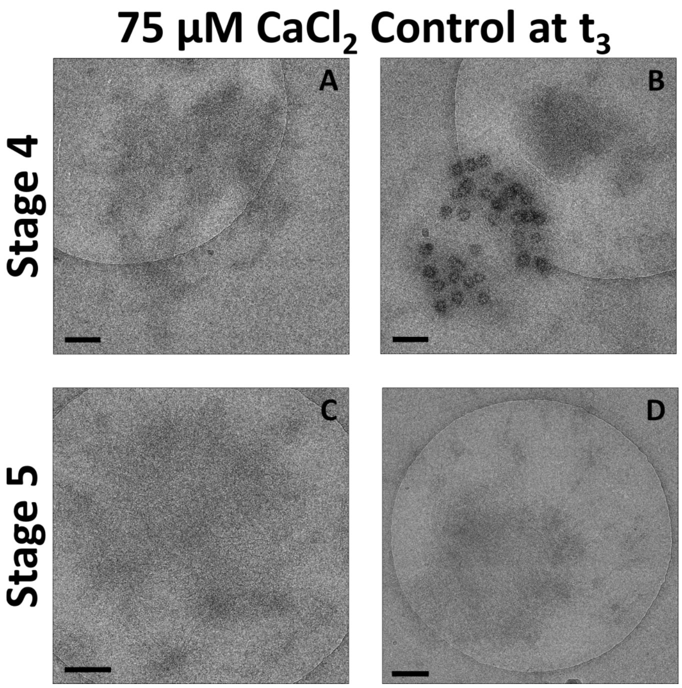


**Figure S4. Massive calcium overload results in severe mitochondrial fragmentation.** Representative images of 10 mins ($t_3$) after the addition of a 75 µM calcium chloride bolus. A-B) Only a few mitochondria with clearly visible granules were observed. C-D) The vast majority of images showed mitochondria that were partially or completely fragmented. Scale bars are 250 nm.

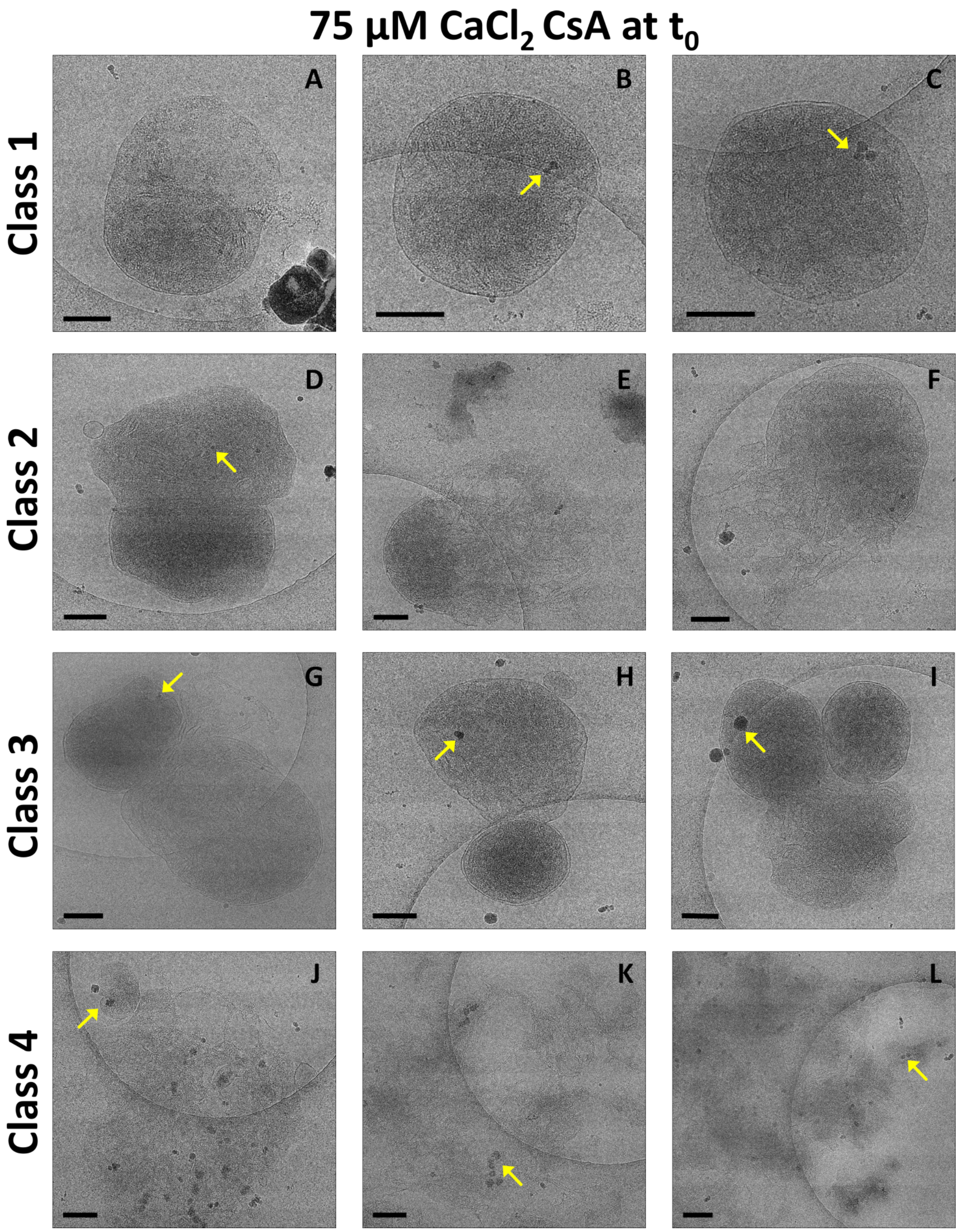


**Figure S5. CsA disrupts the outer membrane morphology, causes release of the inner membrane and tends to form mitochondrial clusters.** Representative images before the addition of a 75 μM calcium bolus ($t_0$) in the presence of 1 μM CsA. Mitochondria were energized with 5 mM sodium pyruvate and 1 mM L-malate. A-C) CsA induced morphological changes in the outer mitochondrial membrane. There are mitochondria with calcium phosphate granules present as shown by the arrows. D-F) While in others the inner membrane is released. G-I) Clusters of mitochondria are commonly observed with treatment. J-L) In some mitochondria, the outer membrane is completely lost. Scale bars are 250 nm.

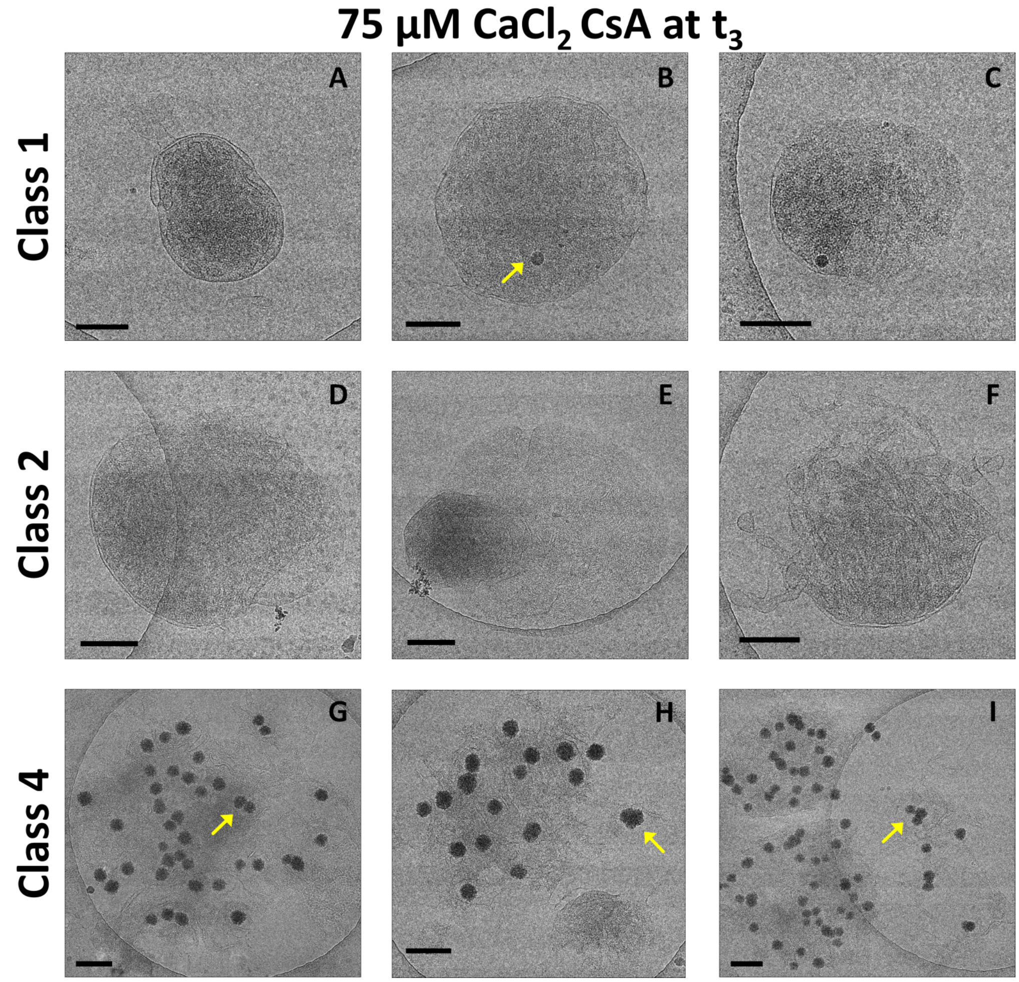


**Figure S6. In the presence of moderate calcium overload, CsA preserves inner mitochondrial membrane intactness.** Representative images 10 mins after the addition of a 75 µM calcium bolus ($t_3$) in the presence of 1 µM CsA. A-C) Mitochondria with or without granules with defined inner and outer membranes. D-F) There are mitochondria with visible cristae ejecting the inner membrane. G-I) After losing the outer membrane, mitochondria spread the inner membrane across the carbon grid. Only 2 micrographs were found in clusters by the time point. However, the presence of mitochondria with unraveled inner membranes in the presence or absence of granules were increased at this time point. Scale bars are 250 nm. Arrows are pointing granules within the mitochondria.

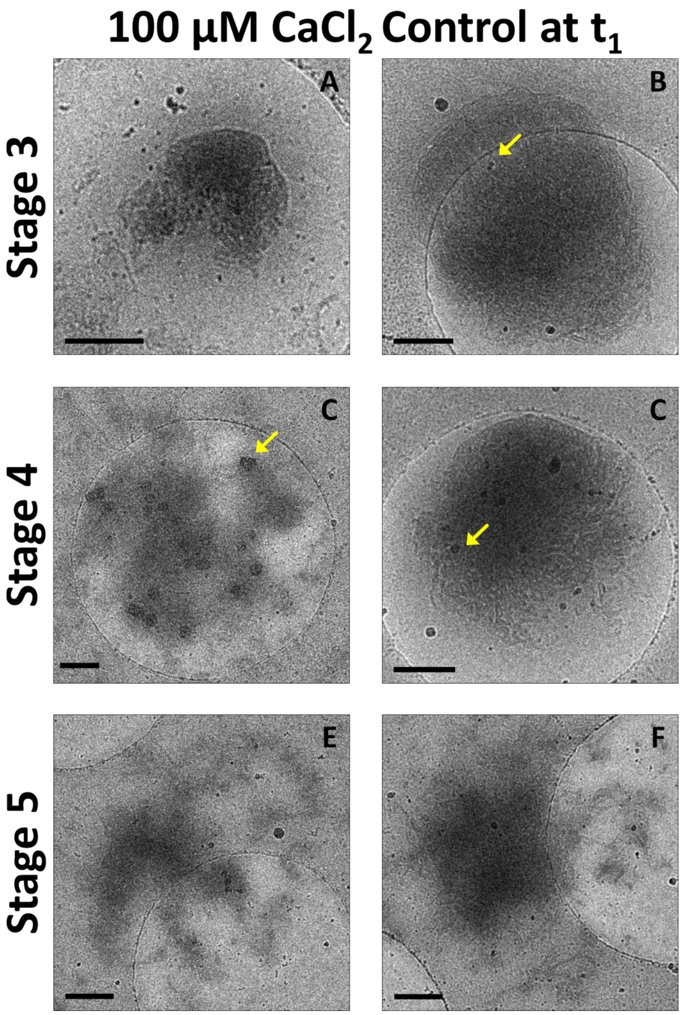


**Figure S7. Mitochondrial morphology is severely compromised and occurs more rapidly at greater calcium loads.** Representative images from isolated mitochondria (0.1 mg/ml) acquired 1.5 mins ($t_1$) after the addition of a 100 µM calcium chloride bolus. Most of the mitochondria were permeabilized under such high calcium load conditions. Only a few stages were identified at this early time point suggesting that membrane fragmentation rapidly occurs. This data is confirmed by the absorbance data where large amplitude swelling was observed. The formation of granules is seen in multiple mitochondria. Scale bars are 250 nm. Arrows are pointing granules within the mitochondria.

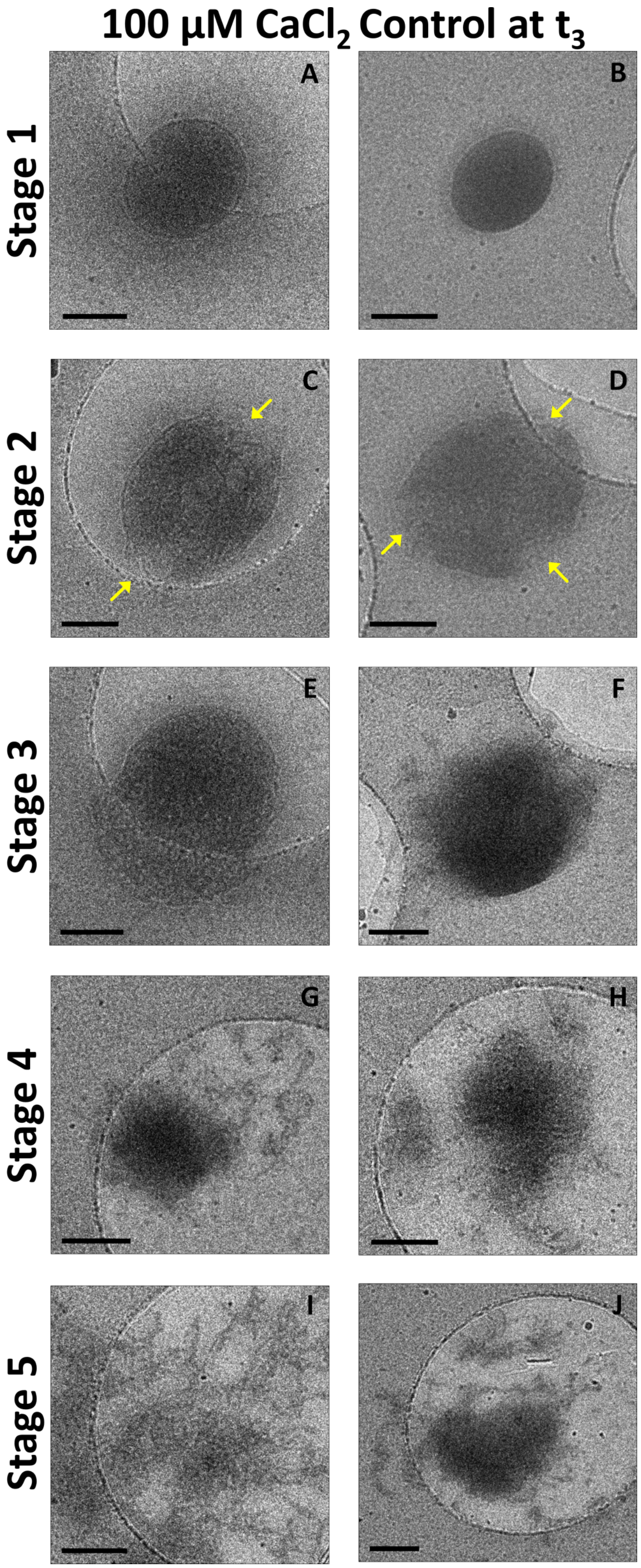


**Figure S8. Heterogeneous responses to high calcium loads leading to mitochondrial permeability transition.** Representative images from isolated mitochondria (0.1 mg/ml) acquired 10 mins ($t_3$) after the addition of a 100 µM calcium chloride bolus. There is a heterogeneous response in the mitochondrial population and multiple stages could be characterized despite being collected at this time-point. Some mitochondria displayed multiple regions of outer membrane rupture as shown by the arrows in stage 2.  Scale bars are 250 nm.

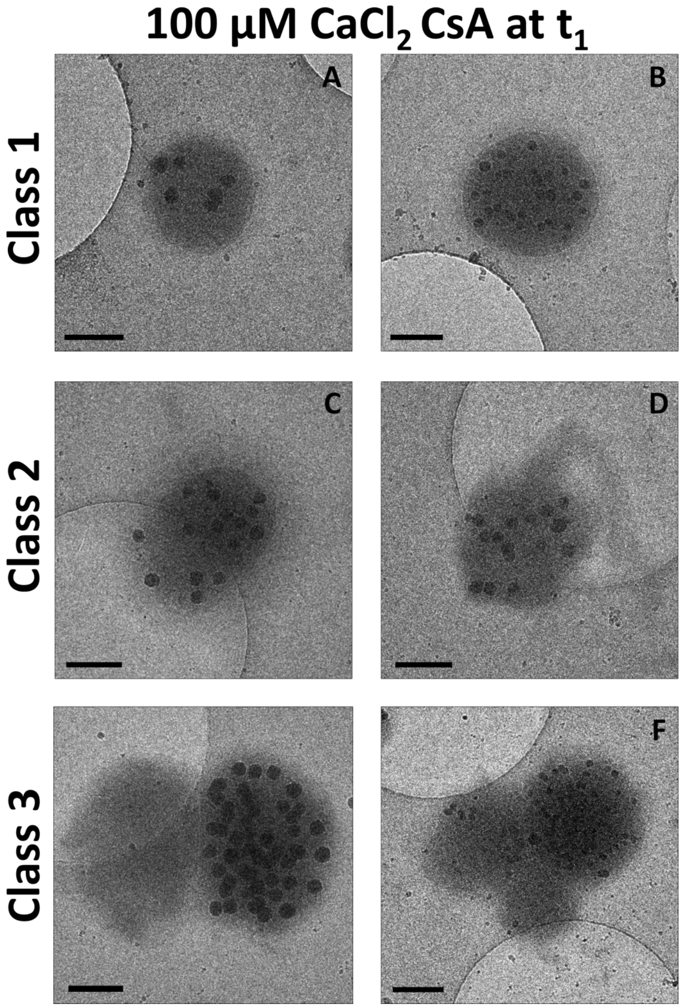


**Figure S9. In extreme calcium overload conditions, CsA preserves inner mitochondrial membrane intactness.** Representative images before the addition of a 100 µM calcium bolus ($t_1$) in the presence of 1 µM CsA. A-B) The presence of calcium led to the formation of granules of various sizes varying between but not within mitochondrion. There is a heterogeneous response to the initial observed effect of CsA on membrane alterations. C-D) The release of inner membrane is apparent in some mitochondrial micrographs. E-F) While others show clusters of mitochondria as observed in the 75 µM calcium chloride bolus experiment. Scale bars are 250 nm.

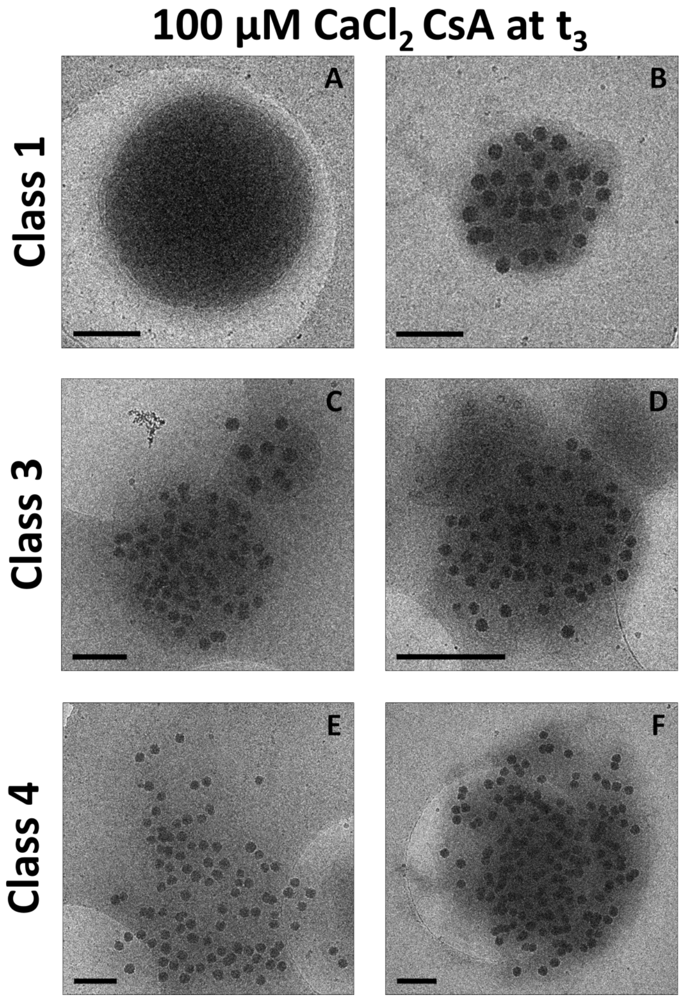


**Figure S10. Mitochondria remain functional at the highest calcium bolus in the presence of CsA.** Representative images from isolated mitochondria (0.1 mg/ml) acquired 10 mins ($t_3$) after the addition of a 100 µM calcium chloride bolus. A-B) There is a heterogeneous response following the addition of a calcium bolus. C-D) Mitochondrial clusters could be identified similar to the previous time points and calcium boluses in the presence of CsA treatment. E-F) Also, mitochondria with protruded inner membranes without outer membranes were detected. Scale bars are 250 nm.

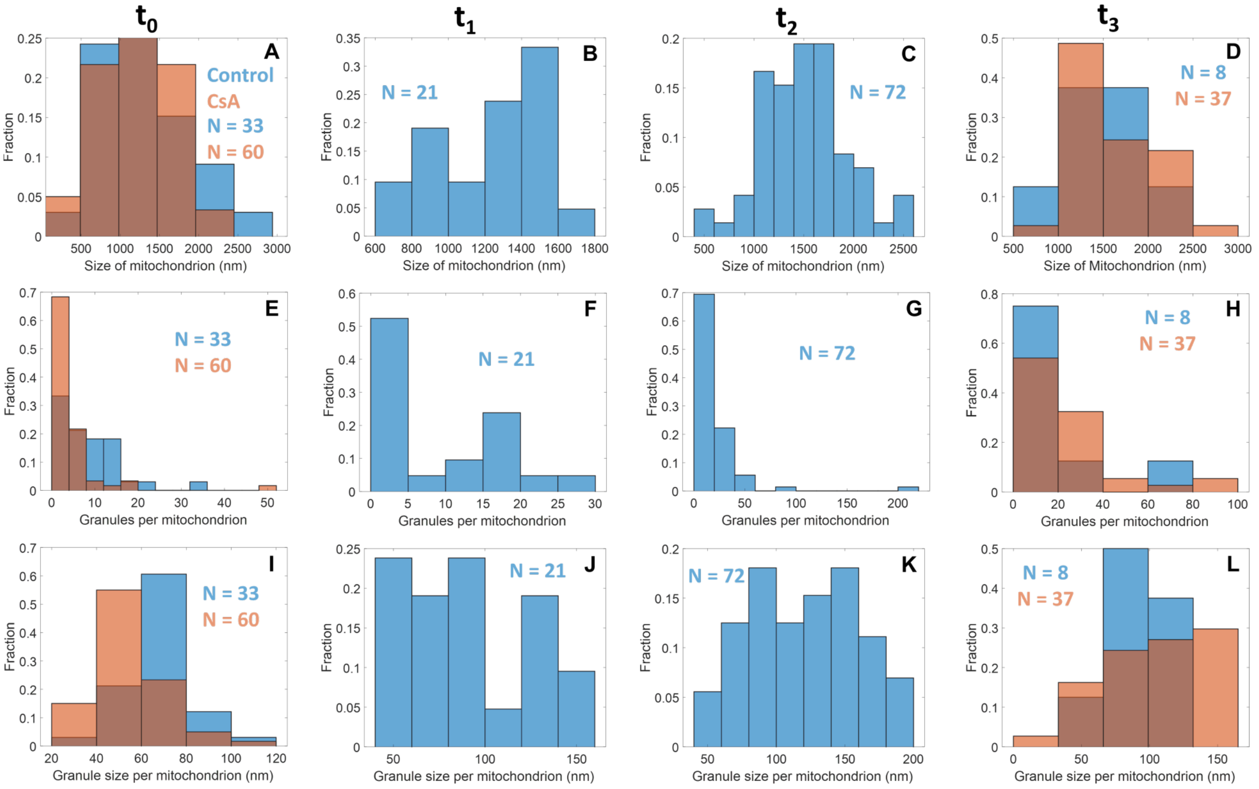


**Figure S11. CsA enhances the number and size of calcium phosphate granules per mitochondrion.** The mitochondrial size, calcium phosphate granules size and number per mitochondrion were quantified for each time-point ($t_0$ – $t_3$) before and after the addition of a 75 μM calcium bolus in the presence or absence of CsA. A) There are no differences in the mitochondrial size before the addition of calcium. D) In contrast, CsA increased marginally their sizes. E and I) Exposing the control mitochondrial suspension to calcium for longer periods increased the number and growth of granules. H and L) However, 10 mins after the addition of calcium ($t_3$) there is a marked decrease in size and number of granules suggesting mitochondrial fragmentation and permeabilization. Conversely, treated mitochondria developed granules of greater size and enhanced the number of granules per mitochondrion.
